# Fast Periodic Stimulation (FPS): A highly effective approach in fMRI brain mapping

**DOI:** 10.1101/135087

**Authors:** Xiaoqing Gao, Francesco Gentile, Bruno Rossion

## Abstract

Functional magnetic resonance imaging (fMRI) is a major technique for human brain mapping. We present a Fast Periodic Stimulation (FPS) fMRI approach, demonstrating its high effectiveness in defining category-selective brain regions. Observers see a dynamic stream of widely variable natural object images alternating at a fast rate (6 images/sec). Every 9 seconds, a short burst of variable face images contrasting with objects in pairs induces an objective 0.111 Hz face-selective neural response in the ventral occipito-temporal cortex and beyond. A model-free Fourier analysis achieves a two-fold increase in signal-to-noise ratio compared to a conventional block-design approach with identical stimuli. Periodicity of category contrast and random variability among images minimize low-level visual confounds while preserving naturalness of the stimuli, leading to the highest values (80-90%) of test-retest reliability yet reported in this area of research. FPS-fMRI opens a new avenue for understanding brain function with low temporal resolution methods.

**Highlights:** FPS-fMRI achieves a two-fold increase in peak SNR over conventional approach

FPS-fMRI reveals comprehensive extended face-selective areas including ATL

FPS-fMRI achieves high specificity by minimizing influence of low-level visual cues

FPS-fMRI achieves very high test-retest reliability (80%-90%) in spatial activation map

**eTOC Blurb:** In Brief

Gao et al. present a novel FPS-fMRI approach, which achieves a two-fold increase in peak signal-to-noise ratio in defining the neural basis of visual categorization while preserving ecological validity, minimizing low-level visual confounds and reaching very high (80%-90%) test-retest reliability.

## Introduction

A fundamental goal of neuroscience is to build a comprehensive map of the human brain, i.e., to define the structure and function of each of its regions (Brodmann, 1909; Amunts & Zilles, 2015; Glasser et al., 2016). The advent of the non-invasive, spatially-resolved functional magnetic resonance imaging (fMRI) technique in the early 1990s (Ogawa et al., 1990; 1992) has provided an unprecedented opportunity to reach this goal, and this technique has now become a major player in Systems and Cognitive Neuroscience. Starting with visual perception, the dominant modality in primates, occupying a substantial fraction of the cortex, researchers have used fMRI to define areas specifically involved in processing low-level visual attributes such as color or motion (e.g., McKeefry & Zeki, 1997; Tootell et al., 1995; Winawer & Withoft, 2015) and retinotopic maps (Sereno et al., 1995; Engel et al., 1997; Wandell & Winawer, 2011). More recently, this approach has been extended to build maps of higher-level areas responding differentially to different categories of the visual world, such as faces, places and body parts (see review by Grill-Spector & Weiner, 2014), and to decode other categories as distributed patterns of variable neural activity across smaller brain volumes (i.e., voxels; e.g., Haxby et al., 2001; Kriegeskorte et al., 2007; Huth et al., 2012).

With a low temporal resolution method such as fMRI measuring neural activity indirectly (i.e., the Blood Oxygenation Level-Dependent, BOLD, response), identifying category-selective brain regions requires presenting visual stimuli belonging to different categories at a relatively *slow* rate, i.e., separated by several seconds, in order to isolate neural activity to each category. Most often though, stimuli from the same condition/category are presented consecutively for 10-20 seconds, i.e., a block-design. The estimated BOLD response during the whole block with respect to a baseline measure (activity to a uniform visual field, or the average activity across all blocks of stimuli) is then considered as reflecting the brain’s response to this category (e.g., Aguirre & D’Esposito, 1999; Weiner & Grill-Spector, 2010). Then, by subtracting neural responses to different categories from one another, category-selective maps, either of regions or patterns of voxels, can be identified (Grill-Spector & Weiner, 2014; Kanwisher, 2017).

Although this approach provides important information regarding human brain cartography, it relies on a number of unwarranted assumptions, such as pure insertion in the context of cognitive subtraction (Friston et al., 1996; D’Esposito, 2010) and uniformity of the modelled hemodynamic response function (HRF) for different brain regions (Boynton et al., 1996; Buxton, 2004). Most importantly, this approach does not consider three key aspects of perceptual categorization when designing stimulation parameters. First, perceptual categorization processes can occur at a high speed in a continuous or quasi-continuous stimulation mode in the visual world (Potter et al., 2012; Retter & Rossion, 2016; Thorpe et al., 1996). This high speed implies that high-level visual areas could be optimally sampled at faster rates than traditionally used in fMRI (Gentile and Rossion, 2014). Critically, the optimal stimulation rate should take into account the duration of a full perceptual categorization process following brief stimulation duration (Retter & Rossion, 2016). Second, to avoid category-specific adaptation (Kovacs et al., 2006) that occurs in block designs when exemplars of the same category are presented consecutively, a direct comparison (i.e., a contrast) between categories should be measured. The final aspect concerns the contribution of low-level visual cues such as differences in global contrast or spatial frequency content to perceptual categorization. Either this issue is not taken into account, using ecological stimuli that leads to large but partially unspecific differential responses (e.g., the “dynamic face localizer approach”, Fox et al., 2009) or, alternatively, stimuli from different categories are normalized for low-level visual cues at the expense of ecological validity (e.g., grayscale segmented full frontal face and house stimuli equalized for power spectra, Rousselet, Husk, Bennett, & Sekuler, 2008). These unwarranted assumptions and suboptimal stimulation parameters add extra noise to the existing physiological, thermal, and scanner noises (Kruger & Glover, 2001), lowering the signal-to-noise ratio (SNR) of fMRI measurement. As a result, extensive trial averaging is often required to achieve effective signal detection (Murphy, Bodurka, & Bandettini, 2007). For these reasons, fMRI studies in cognitive neuroscience may suffer from low sensitivity, specificity and reliability (Bennet & Miller, 2010).

Here, we take into account all these aspects to introduce an extremely effective approach to localize category-selective neural responses. We use this approach as a model to measure brain function with fMRI or low temporal resolution methods in general. In this approach, visual stimuli are presented at a *fast* presentation rate (6 images/sec, 6Hz) allowing one fixation by image – largely sufficient for perceptual categorization (e.g., Potter et al., 2012; Retter & Rossion, 2016; Thorpe et al., 1996) – throughout the entire recording of neural activity (Figure 1a; Supplemental Movie S1). This dynamic stimulation sets a high baseline level of activity in low-level visual areas as well as in non-category-selective high-level visual areas. Then, we introduce transient switches from non-target object categories to a target category, here faces (Figure 1a, red bins). In populations of neurons responding selectively to faces, such transient switches elicit differential neural responses that *directly* reflect category-selectivity because they contrast with the continuous stimulation stream of non-face objects. Hence, contrast is maximized, and inference regarding category-selectivity can be made without post-hoc cognitive subtraction. The transient switches to faces are grouped in short (2secs) bursts, to further improve SNR in fMRI. However, within a burst, faces appear only every two stimuli, i.e. alternate with a randomly selected object (Figure 1b). This way, category-specific adaptation (e.g., Kovacs et al., 2006) is reduced during the burst, multiple contrasts are measured within a burst, and the temporal separation between two faces is more than 300ms, leaving sufficient time for occurrence of the bulk of the face-selective neural response as estimated with scalp EEG recording (Retter & Rossion, 2016). Critically, we set the bursts to appear at a *fixed* frequency, so that the magnitude of the differential neural responses can be measured without HRF modelling, i.e., as the Fourier amplitude of the frequency of the switches (e.g., Bandettini et al., 1993; Engel, Glover, & Wandell, 1997; Puce et al., 1995). This model-free approach therefore allows fair comparisons across brain regions and individuals. It has high SNR, since it is only affected by noise occurring at the exact same frequency of the switches, not broadband frequency noise (Regan, 1989). Finally, a wide variety of natural images, which have complex statistical properties (Simoncelli & Olshausen, 2001), ensure that the succeeding images represent many different types of low-level contrasts, minimizing the contribution of specific low-level visual cues to category-selective responses occurring at the same periodic frequency (Rossion et al., 2015). At the same time, widely variable images of faces are used to ensure that a category-selective response is not tied to specific exemplars (i.e., is generalized). Based on these unique features, we name the current approach as a Fast Periodic Stimulation (FPS)-fMRI paradigm.

**Figure 1.**
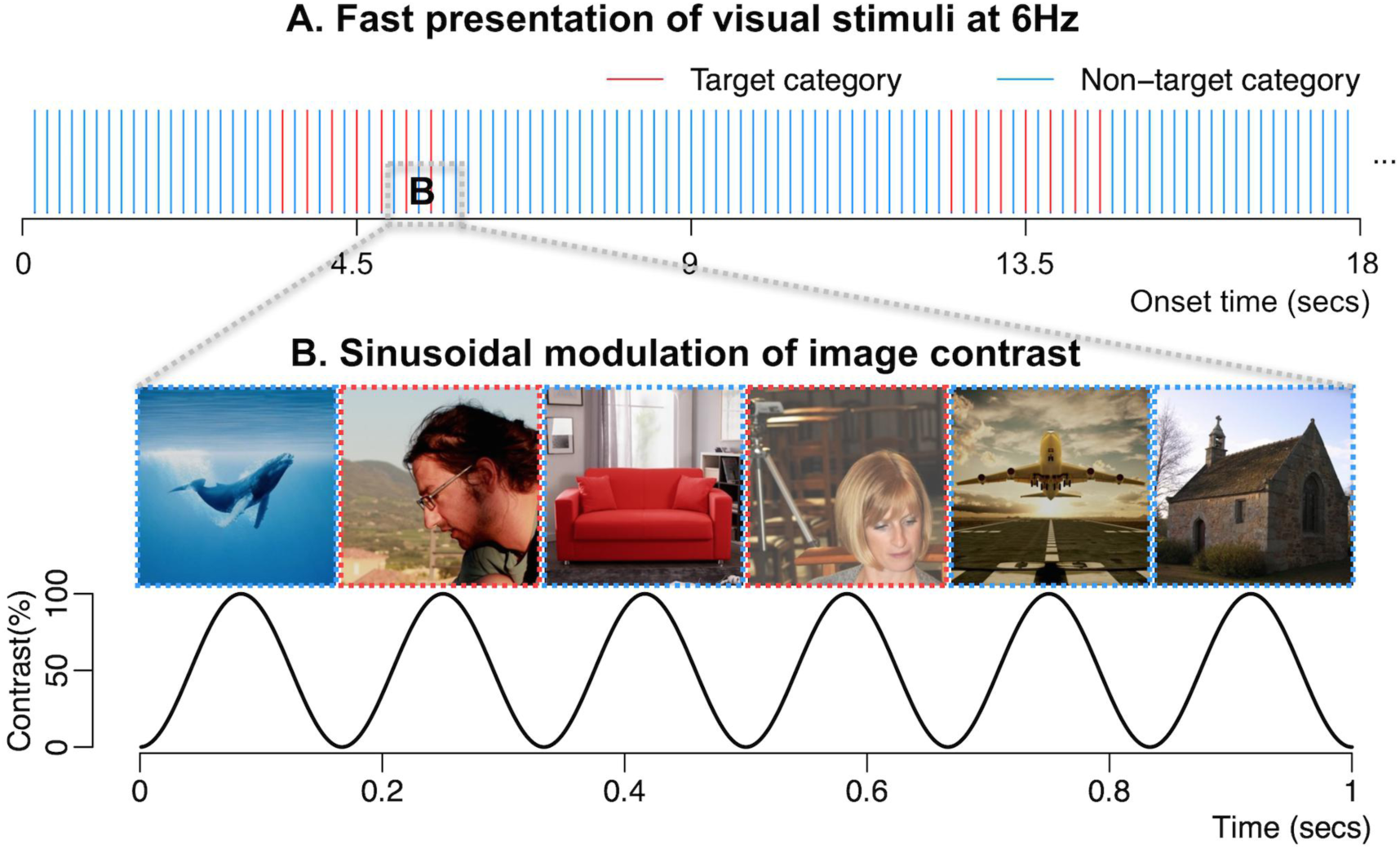
The Fast Periodic Stimulation (FPS)-fMRI paradigm. (A) Stimuli from non-target categories alternate at a rapid rate (6 Hz, blue bins, see Supplemental Movie S1). Every 9 seconds (i.e., 54 stimuli), a “burst” of stimuli from the target category (faces, red bins) is presented for 2.167 seconds. This face burst contains 7 faces alternating with 6 non-face objects to form direct contrasts between faces and objects. Only sections of the sequences are shown in the figure. (B) Example face and object alternations (see Supplemental Fig. S1 for more example images) and image contrast modulation (0% to 100%).

We designed a functional “face localizer” based on the above-mentioned principles. After Kanwisher and colleagues (1997), face localizers are arguably the most widely used approach to define category-selectivity in neuroimaging, both in human and non-human primates (Tsao et al., 2008; Kanwisher, 2017). In humans, the representation of faces differs from other object categories at the level of a large number of distributed regions, or functional clusters, which have been reported by numerous studies primarily in the ventral occipito-temporal cortex (VOTC), but also in specific regions of the anterior temporal lobe (ATL), superior temporal sulcus (STS), and to a lesser extent in the parietal and frontal lobes (Haxby et al., 2000; Duchaine & Yovel, 2015). Therefore, it provides an excellent model to assess the validity of the novel FPS-fMRI approach. For comparison, we also ran a conventional face localizer task based on a block-design with the exact same stimuli and same duration of scanning.

We demonstrate that FPS-fMRI: (1) objectively identifies the well-known core face-selective areas and their right hemispheric dominance with extremely high SNR in all individual brains; (2) substantially increases the detection of consistent face-selective clusters in higher order brain regions such as the ATL despite magnetic susceptibility artefacts; (3) eliminates the contribution of low-level visual areas, i.e. the primary visual cortex to category-selective brain responses while preserving naturalness of the images; (4) achieves the highest test-retest reliability ever reported in imaging higher level brain regions.

## Results

Twelve participants each performed three runs of the FPS face localizer task (FPS-face), three runs of the conventional face localizer task (CONV-face), and two runs of FPS task with scrambled images (FPS-scrambled). In the FPS-face condition, a continuous stream of non-face images was presented at a fast rate (6Hz). Every 9 seconds (0.111Hz, referred to as face stimulation frequency), a short burst of faces contrasting with objects appears for 2.167 seconds. The CONV-face condition was based on a block-design with alternating 18-secs face blocks and non-face object blocks. The FPS-scramble condition had the same sequence as the FPS-face condition but with Fourier phase scrambled versions of face and object images. All the runs had the same duration (396 secs).

### FPS-fMRI effectively defines category-selective BOLD responses

To illustrate the effectiveness of the FPS-fMRI paradigm in defining category-selective BOLD responses, we first consider a representative voxel, identified by showing the largest faces > objects contrast in a single brain in the conventional face localizer (Figure 2a). This voxel is located in the area showing the largest face-selective response in the human brain, the lateral section of the right fusiform gyrus, often defined as the *Fusiform Face Area* (FFA, Kanwisher et al., 1997). Fast Fourier Transform (FFT) applied to the BOLD response time course of this voxel (Figure 2b) reveals a high signal at the face stimulation frequency (0.111 Hz) in the FPS-face condition (Figure 2c). Given the very high frequency resolution (1/396secs = 0.0025 Hz), the signal is concentrated on a tiny frequency bin in the amplitude spectrum of the BOLD response. For this example voxel, all of the response of interest concentrates on the fundamental frequency, i.e., 0.111 Hz. In the same region in a few other individual brains, there were also negligible amplitude increases at the second harmonic (0.222 Hz) (Supplemental Fig. S2).

**Figure 2.**
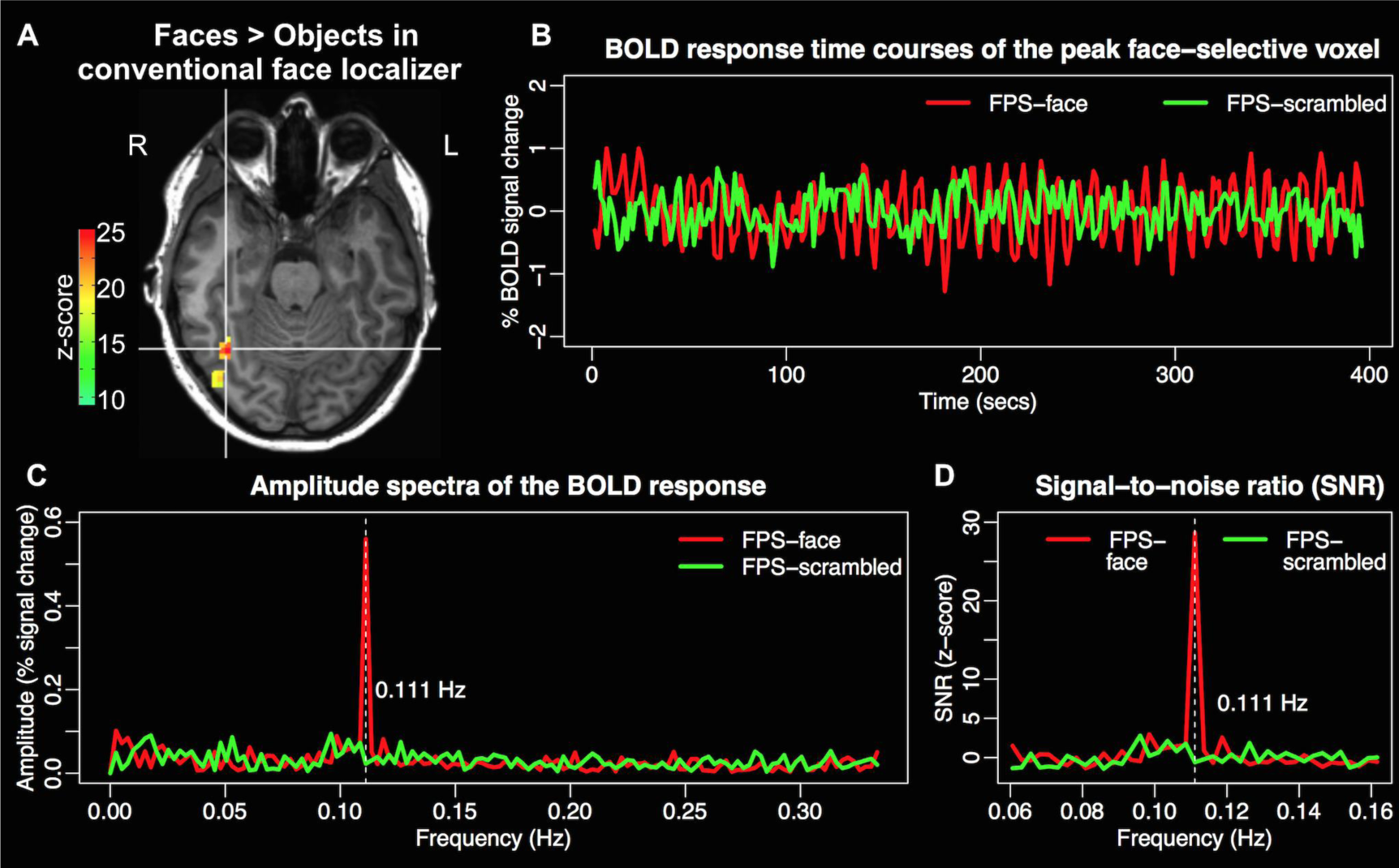
BOLD response in an example face-selective voxel. (A) Peak face-selective voxel (cross hair, right lateral fusiform gyrus) defined by faces > objects contrast in the conventional face localizer. (B) BOLD response time courses of the peak face-selective voxel in the FPS paradigm with natural images (FPS-face, in red) or with phase-scrambled images (FPS-scrambled, in green). (C) Amplitude spectra of the BOLD response. The white dashed line indicates the face stimulation frequency (0.111 Hz). (D) Signal-to-noise ratio (z-score) of the amplitudes.

We calculated a z-score of the amplitude of BOLD response at the face stimulation frequency relative to the noise level in the neighboring frequencies for each voxel as done in previous frequency-based fMRI analysis (McCarthy et al., 1994; Puce et al., 1995). By definition, a z-score is the baseline corrected signal level over noise level measured as the noise standard deviation, and is thus considered as a measure of SNR of the face-selective neural activity (Welvaert & Rosseel, 2013, Figure 2d). In all individual brains, the FPS-face condition achieved very high SNR (z-score ranged from 13.9 to 31.0, mean = 20.8 ± 5.2) at the face stimulation frequency for the peak face-selective voxel in the right FFA identified by the conventional face localizer. Therefore, the FPS-fMRI paradigm can effectively modulate the BOLD signal representing a differential neural response to faces *vs.* non-face objects. Importantly, there was no signal at the face stimulation frequency for FPS-scrambled condition in any of these voxels (z-score mean = 0.25 ± 1.04, *t*(11)=0.83, *p* = 0.42, two-tailed test against 0). The absence of response in the BOLD amplitude spectrum of the same voxels for phase-scrambled images implies that low-level image properties contained in the power spectrum did not contribute to the peak of the face-selective BOLD response at the face stimulation frequency in the right FFA, as well as in other face-selective regions (Supplemental Fig. S3).

### Mapping face-selective cortical areas (group analysis)

To map face-selective brain areas, for each voxel in each individual brain, we calculated a SNR (z-score) of face-selective neural response in the FPS-face condition. For the conventional face localizer (CONV-face), we obtained a z-score from faces > objects contrast for each voxel in each individual brain. We then averaged the z-maps across individual brains using a surface-based averaging approach to identify face-selective areas that are present consistently across participants.

In the FPS-face condition, as shown in Figure 3a, with a conservative threshold level of uncorrected *p* < 10^-6^ (equivalent to a Bonferroni corrected *p* < 0.05), in both hemispheres, we identified face-selective areas in the well-known core face processing network (Haxby et al., 2000; Duchaine & Yovel, 2015) consisting of three cortical areas, namely the “fusiform face area (FFA)” in the middle section of the lateral fusiform gyrus (FG), the “occipital face area (OFA)” in the inferior occipital gyrus (IOG), and the posterior superior temporal sulcus (pSTS). In both hemispheres, the most significant response, i.e. peak of face-selectivity, corresponds to the FFA, and the highest average z-score is found in the right FFA, with a typical 3D coordinate in a normalized brain (42, -54, -14 in Talairach coordinates, compare, for instance, to the right FFA coordinate in Kanwisher et al., 1997: 40, -55, -10; Zhen et al., 2015, for pFFA: 42, -51, -14 (converted from MNI to Talairach); Jonas et al., 2016, FFA identified in intracerebral recording: 41, - 45, -16).

**Figure 3.**
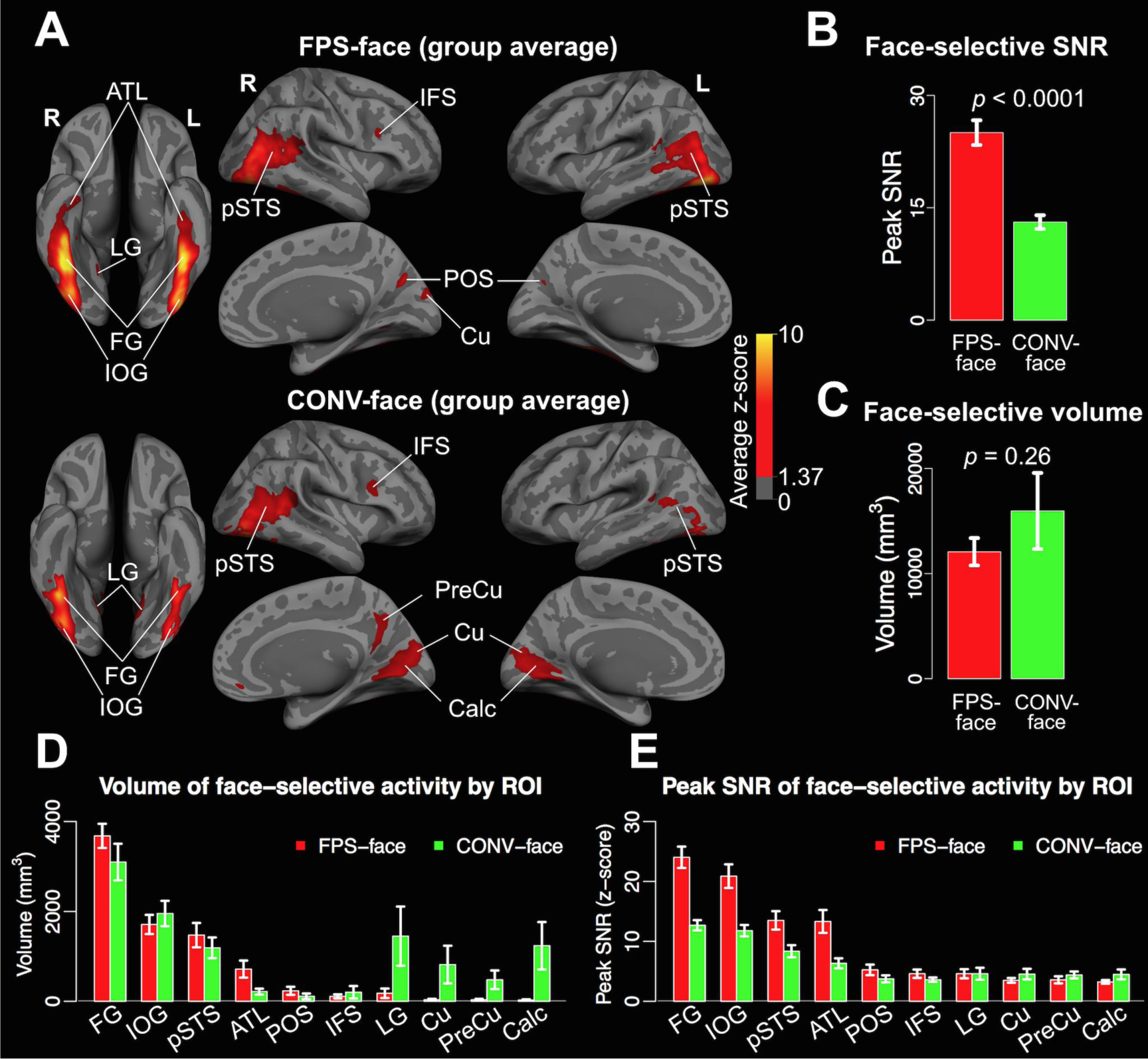
Face-selective activity in the whole cortex and in anatomic ROIs. (A) Face-selective cortical areas identified based on group averaged z-maps (*n* = 12). The maps are thresholded at *p* < 10^-6^, uncorrected, and are projected onto an inflated average cortical surface of all the participants. (B) Peak SNR of face-selective activity in the whole cortex in the FPS-face condition and CONV-face condition, averaged across participants. Data are represented as mean ± SEM. (C) Total volume of face-selective responses in the whole cortex averaged over individual brains with a threshold level of *p* < 10^-4^, uncorrected. Data are represented as mean ± SEM. (D) Volume of face-selective activity in anatomical ROIs averaged over individual brains with a threshold level of *p* < 10^-4^, uncorrected. Data are represented as mean ± SEM. (E) Peak SNR of face-selective activity in anatomical ROIs, averaged across participants. Data are represented as mean ± SEM. *Note: in (D) and (E) we combined homologous ROIs across hemispheres, as the results were similar for both hemispheres. FG: fusiform gyrus; IOG: inferior occipital gyrus; pSTS: posterior superior temporal sulcus; ATL: anterior temporal lobe; POS: parieto-occipital sulcus; IFS: inferior frontal sulcus; LG: lingual gyrus; Cu: cuneus; PreCu: precuneus; Calc: calcarine sulcus.*

Importantly, the FPS-face condition also revealed consistent face-selective activity in the anterior temporal lobe (ATL) in both hemispheres (Figure 3a). The role of ATL in face-selective processing is known from intracerebral recordings (Puce et al., 1999; Jonas et al., 2016) and human lesion studies causing individual face recognition impairments (e.g., Busigny et al., 2014) but has only been recently brought to attention in fMRI (Rajimehr et al., 2009; Nasr & Tootell, 2012; see review by Collins & Olson, 2014), due to the difficulty in identifying these regions with magnetic susceptibility artifacts (Axelrod & Yovel, 2013; Jonas et al., 2015) (see Supplemental Fig. S4 for an example of MRI signal drop in the ATL).

Besides the core face processing network and the ATL, the FPS-face condition also identified consistent face-selective activity in areas previously reported to be involved in face processing in neuroimaging studies. These areas include the Inferior Frontal Sulcus (IFS; e.g., Fox et al., 2009; Ishai et al., 2005; Chan & Downing, 2011), Lingual Gyrus (LG; e.g., Gobbini & Haxby, 2006; Rossion et al., 2012), Cuneus (e.g., Benuzzi et al., 2007; Rossion et al., 2012), and Parieto-Occipital Sulcus (POS; e.g., Tuladhar et al., 2007; Jokish & Jensen, 2007).

In the CONV-face condition, at the same threshold level, we also identified the core face-selective network including bilateral FFA, OFA, and pSTS. However, we noticed three marked differences in comparison to the FPS-face condition: 1) the magnitude of face-selective activation in the FPS-face condition is much higher than in the CONV-face condition. The average maximum SNR (z-score) in the FPS-face condition (25.0±5.7) is about two-fold higher than in the CONV-face condition (13.1±3.2, *t*(11) = 6.2, *p* < 0.001, *Cohen’s d* = 2.57, Figure 3b). Notably, such an increase in the peak face-selective neural responses in the FPS approach compared to the conventional approach is found in every individual brain tested (range of the ratio of increase: 1.1 to 3.4). At the same time, there is no significant difference between the total volumes of face-selective activity identified in the two conditions (*p* = 0.26, Figure 3c); 2) at this threshold level (*p* < 10^-6^, uncorrected), the CONV-face condition failed to reveal face-selective activity in ATL in the group averaged map (Figure 3a); 3) we found extensive activity in the low-level visual regions (e.g., Calcarine sulcus, Figure 3a and 3d) only in the CONV-face condition.

### Face-selective cortical areas in individual brains

Complementary to the group activation maps, we quantified face-selective neural activity in each individual brain with a threshold level of *p* < 10^-4^, uncorrected. This is a necessary step because substantial variability exists in brain structure as well as in functional localization (Frost and Goebel, 2012; Rossion et al., 2012; Zhen et al., 2015; Zilles and Amunts, 2013;). As can be seen in Figure 4, the face-selective activations identified in the FPS-face condition and the CONV-face condition are more focal in individual brains than in the averaged maps (Figure 3a).

**Figure 4.**
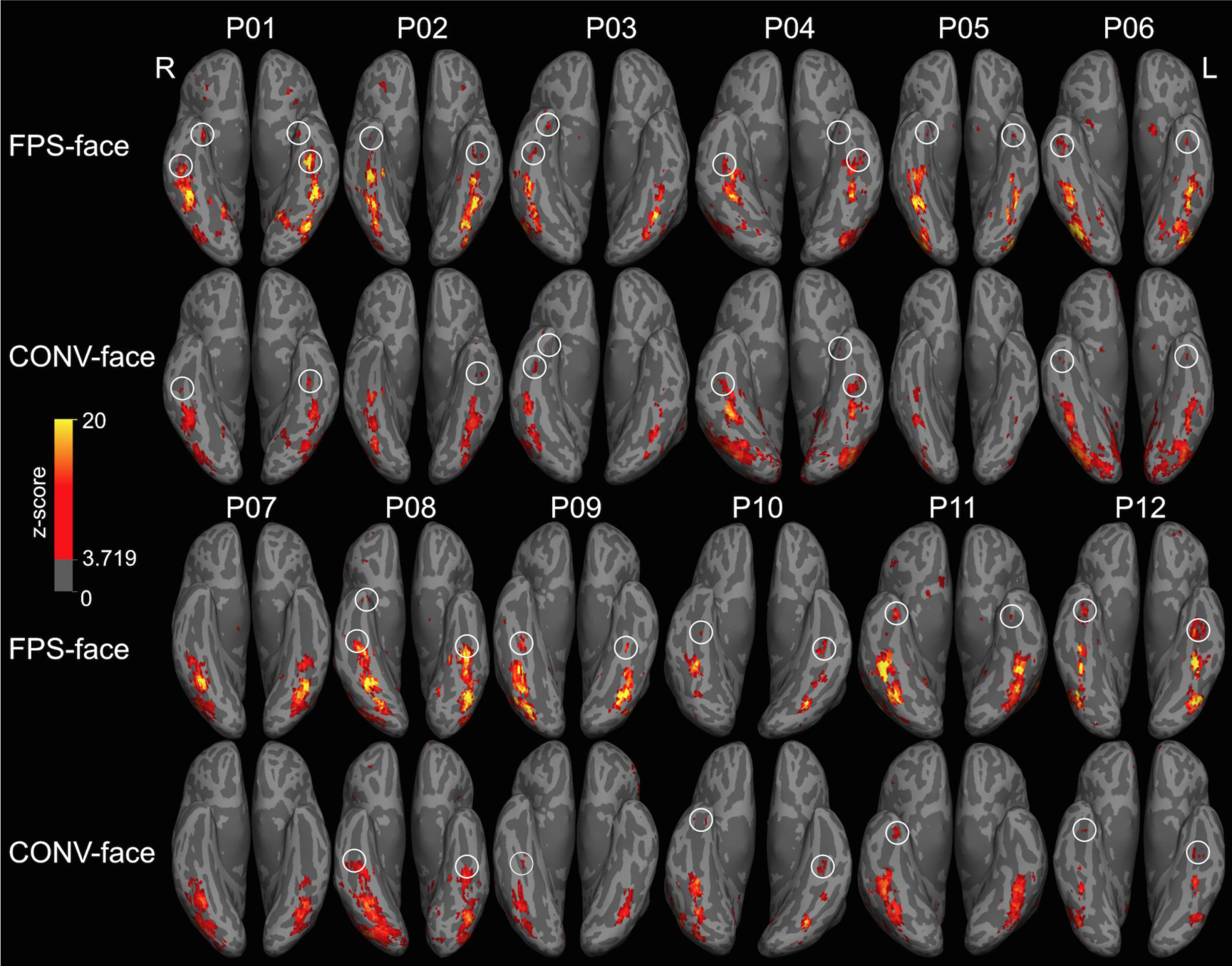
Individual maps of face-selective activity in VOTC. Magnitude (z-score) of face-selective activity was projected to an inflated cortical surface of each individual brain for the FPS-face and the CONV-face conditions, with a threshold level of *p* < 10^-4^, uncorrected. Note the overlap between the face-selective regions identified by the two conditions, but with much higher magnitude reached in the FPS paradigm, leading to extra activation clusters in the anterior temporal lobe (highlighted with white circles). See Supplemental Fig. S5 for lateral and medial views of face-selective activity in all the individual brains.

In both paradigms, consistent with previous neuroimaging studies (e.g., Kanwisher et al., 1997; McCarthy, Puce, Gore, & Allison, 1997; Sergent et al., 1992, Rossion et al., 2012; Zhen et al., 2015), we found larger volume of face-selective activity in the right than in the left hemisphere (for the whole cortex, *p* = 0.0001 for FPS-face condition; *p* = 0.001 for CONV-face condition; the right hemisphere advantage is also significant within ROIs: FG (*p* = 0.0001), pSTS (*p* = 0.004), and POS (*p* = 0.009) for the FPS-face condition; FG (*p* = 0.0003), pSTS (*p* = 0.001), and IOG (*p* = 0.04) for the CONV-face condition), and the location associated with high response magnitude is fairly consistent across the two paradigms for all individual brains (Figure 4).

The overlap between the two paradigms accounted for 59.7± 26.5% of face-selective volume in the CONV-face condition. The percentage was slightly higher 61.3 ± 17.2 % (but not statistically different, *p* = 0.9) for the FPS-face condition. While the percentage of brain activation overlap may appear relatively low in regard to the visual similarity between brain maps in the two conditions (e.g., Figure 4), in a single high-level region, e.g. the right FG, overlap between the two conditions accounted for a higher percentage (90% for CONV-face and 73% for FPS-face, *p* = 0.06). Hence, 90% of face-selective volume in the right FG identified by the CONV-face condition was also identified by the FPS-face condition. Where are the voxels that are activated specifically in each condition? To answer this question, we quantified the volume of above-threshold voxels in 10 anatomically defined brain areas (combining homologous areas across hemispheres) selected based on results of the group analysis. As shown on in Figure 3d, while the highest proportion of volume activated was found in the three core regions of the face network (FG, IOG, pSTS) plus ATL in the FPS condition, this proportion was slightly lower in the CONV-face condition.

Critically, the volume of face-selective area in ATL was significantly larger in the FPS-face condition than in the CONV-face condition (*p* = 0.011). This is particularly important in the context of the relatively recent focus on ATL in face processing and the general difficulty in identifying these regions in fMRI due to magnetic susceptibility artifacts. Considering individual brains, the FPS approach was able to identify regions in 11 (Figure 4) individual brains out of a total of 12, without using specific fMRI sequences to reduce magnetic susceptibility artifacts in the temporal lobe (Devlin et al., 2000).

In contrast, a much larger volume of “face-selective” area was found in low-level visual regions such as the Calcarine sulcus in the CONV-face condition than in the FPS-face condition in both hemispheres (*p* = 0.039, see Figure 3d and Figure 5). This finding suggests that the conventional face localizer is more affected by low-level visual differences between faces and non-face object categories, as predicted.

**Figure 5.**
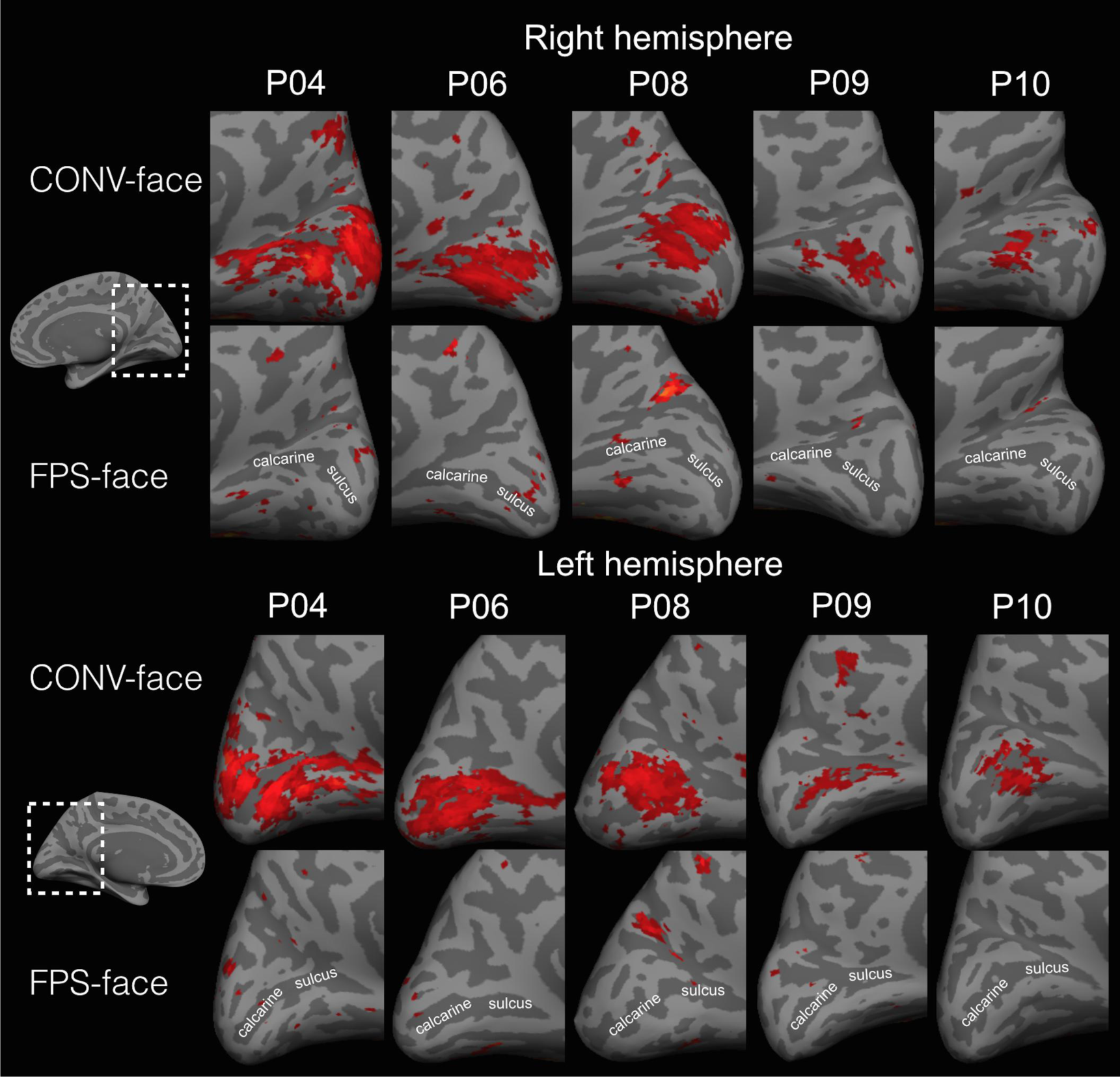
“Face-selective” activity in Calcarine sulcus. Inflated surfaces of the medial occipital cortex of five example participants show “face-selective” activity in bilateral Calcarine sulcus in the CONV-face condition but not in the FPS-face condition. The maps are thresholded at *p* < 10^-4^, uncorrected.

Besides volume, the FPS paradigm achieved much higher face-selective response magnitude in higher-level areas than the CONV-face condition (Figure 3e; FG, *p* = 0.0001; IOG, *p* = 0.001; pSTS, *p* = 0.008; ATL, *p* = 0.0005).

In summary, the FPS approach is much more sensitive to detect face-selective neural activity in typical high-level visual areas and the anterior temporal lobe, while being less contaminated by low-level confounds (i.e., more specific) than a conventional localizer procedure in fMRI.

### Test-retest reliability

How reliable, or replicable, are these observations, is a key issue in the current fMRI research (Bennett and Miller 2010; Nichols et al., 2016). To assess the reliability of the FPS paradigm, we analysed data of each individual run in the FPS-face condition. In comparison, we also analysed the individual runs in the CONV-face condition. Figure 6a provides a visual illustration of the consistency across runs in one representative participant: there is a good correspondence of face-selective voxels across different runs within both the FPS-face condition and the CONV-face condition.

**Figure 6.**
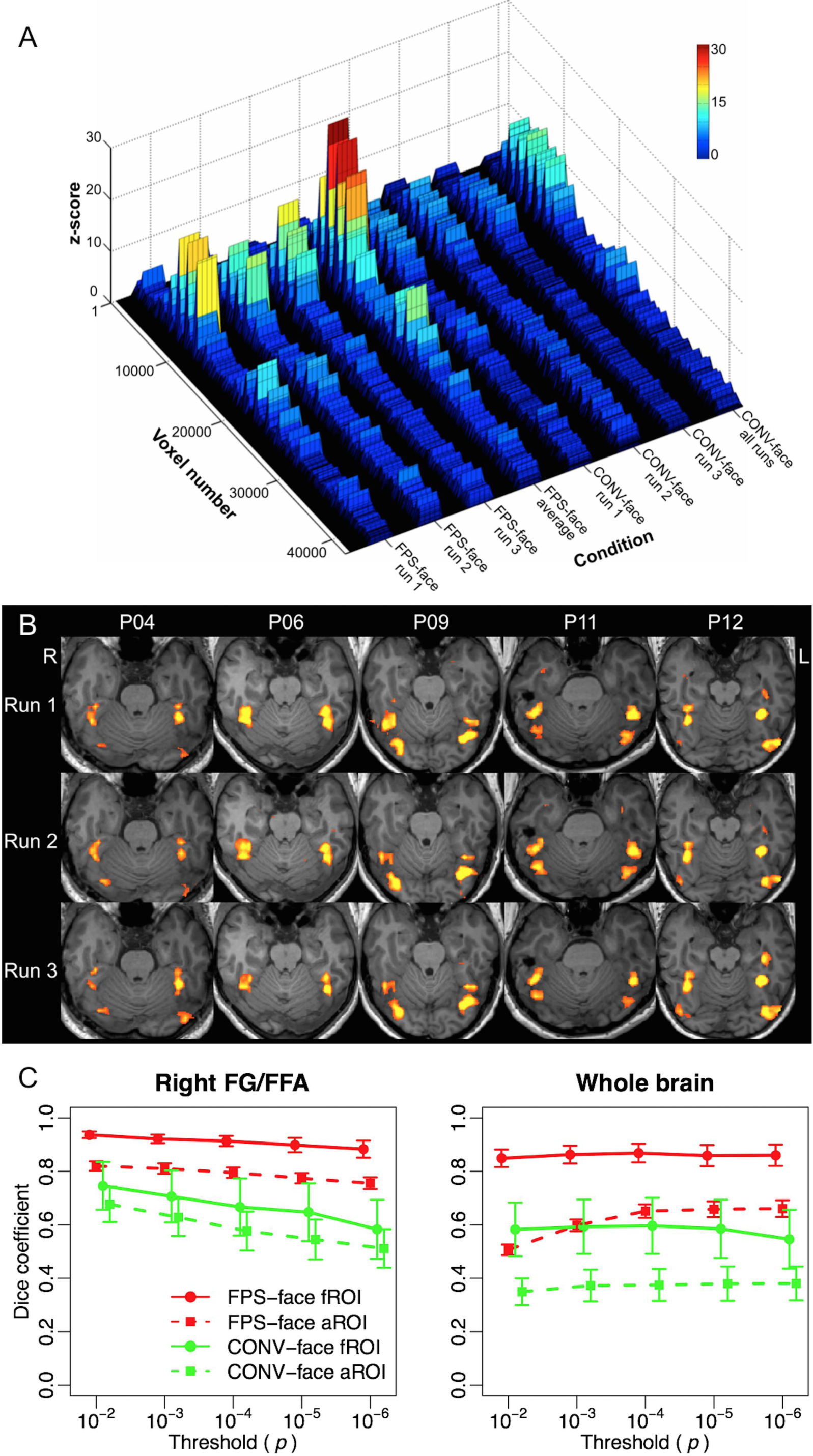
Test-retest reliability of face-selective spatial activation maps. (A) An example of face-selective neural response in each run of the FPS-face condition (FPS-face runs 1, 2, 3) and the CONV-face condition (CONV-face runs 1, 2, 3) in one participant (see Supplemental Fig. S6 for all the participants). “FPS-face run average” was calculated based on the average BOLD response time course of the three FPS-face runs. “CONV-face all runs” was calculated based on data from all the three CONV-face runs. Voxels in the whole brain were listed on a one-dimensional axis (Voxel number), so only spatial correspondence between different conditions was maintained. (B) Example spatial activation maps of three FPS-face runs in five participants (threshold level, *p* < 10^-4^, uncorrected). (C) Dice coefficient of spatial overlap of super-threshold voxels across three runs in the FPS-face and CONV-face conditions at five levels of thresholds. Data are represented as mean ± SEM. Left panel: reliability analyses confined within anatomically defined right fusiform gyrus (aROI) or the functionally defined right FFA based on the first run (fROI). Right panel: reliability analyses in the whole cortex (aROI) or within the super-threshold voxels based on the first run (fROI).

To quantify the overlap between face-selective voxels identified between different runs, we calculated a consistency score using Dice coefficient (see methods). First, we confined the calculations within an anatomically defined area containing only voxels in the right fusiform gyrus (the anatomical ROI or aROI approach) to make the current measure comparable to values reported in previous studies (Berman et al., 2010; Duncan et al., 2009, 2011). At a threshold level of uncorrected *p* < 10^-4^, the face-selective voxels identified in different runs in the CONV-face condition reached an average value of 0.58 ± 0.25, which is in the high end of the typical range as reported in previous studies (Berman et al., 2010; Duncan et al., 2009; Duncan & Devlin, 2011). However, the FPS-face condition reached a significantly higher consistency (0.80 ± 0.07, range = 0.67 to 0.90, (*t*(11) = 3.08, *p* =0.01), reaching 80% overlap of face-selective voxels between runs on average. Note that this consistency value is obtained in a ROI, the right fusiform gyrus, containing about 30% of voxels above threshold, so that brain activity in the vast majority of voxels may vary randomly from run to run and decrease this consistency value. Another approach is to confine the analysis to the right FFA in every individual as defined on the first run of each condition (the functional ROI, or fROI approach), and calculate the Dice index using data from the other two runs. Here, consistency values reach 0.91±0.07 (0.79 to 0.99, see Figure 6b for a visual demonstration of consistency in the FFA in individual participants) in the FPS-face condition, significantly higher than in the CONV-face condition (0.67 ± 0.37; *t*(11)= 2.3, *p* = 0.04).

Since test-retest reliability may decrease with increasing threshold (Berman et al., 2010; Duncan et al., 2009; Duncan & Devlin, 2011), we calculated the Dice coefficients at five levels from a liberal (uncorrected *p* < 10^-2^) to a conservative (uncorrected *p* < 10^-6^, equivalent to a Bonferroni corrected *p* < 0.05) threshold. We performed the calculation separately within the anatomically defined right fusiform gyrus and the whole cortex (aROI approach). We also calculated the reliability score within functionally defined right FFA based on the first run of each condition, and all the super-threshold voxels in the whole cortex in the first run (fROI approach).

In the CONV-face condition, reliability decreased with increasing threshold (Figure 6c) in both the anatomically defined right fusiform gyrus (slope = -0.04, 95%CI = -0.054 to -0.030) and the functionally defined right FFA (slope = -0.04, 95%CI = -0.067 to -0.019), as previously reported (Berman et al., 2010; Duncan et al., 2009; Duncan & Devlin, 2011). However, the reliability decreased less steeply with increasing threshold in the FPS-face condition (slope = -0.016 for aROI approach and -0.012 for fROI approach), achieving Dice coefficients of around 0.9 at all threshold levels in the right FFA. To the best of our knowledge, this is the highest test-retest reliability of spatial overlap of activation maps across fMRI scanning runs yet reported in a high-level functional area.

In the whole cortex (Figure 6c), reliability was lower in general compared to corresponding values from the right fusiform gyrus (aROI) or right FFA (fROI). Nevertheless, in the FPS-face condition, within a functional mask defined by the super-threshold voxels from the first run, the reliability score reached above .8 at all threshold levels. In contrast, the highest average reliability score in the CONV-face condition was below .6.

In summary, while the conventional face localizer used in this study is at least as valid and reliable as those used in previous studies, the FPS-fMRI approach still greatly enhances test-retest reliability of category-selective neural activation.

### The respective contribution of FFT analysis and stimulation mode

Two main factors can contribute to the power of the FPS-fMRI paradigm. First is the fast-paced visual stimulation, and the direct contrast between faces and non-face categories comparing to a slow-paced, blocked conventional face localizer in which category-specific adaptation effects could occur. Second is the data analysis procedure. In this regard, FFT analysis is only affected by narrow-band noise in the same frequency as the face stimulation (Regan, 1989). In contrast, the General Linear Model (GLM) approach is affected by broadband noise. In addition, the data analysis approach is model-free in FPS-fMRI, while the GLM approach is based on assumptions of HRF function. We further investigated the respective contribution from these two aspects by (1) reanalyzing data of the CONV-face condition with FFT (since we altered face blocks and non-face blocks at a fixed interval in that condition) and reanalyzing data from the FPS-face condition using the GLM approach by modelling each face burst as an event. We investigated the respective contribution of *stimulation paradigm* and *analysis procedure* to the maximum face-selective response in three face-selective areas showing maximal differences between the FPS-face condition and CONV-face condition: the FG, ATL, and Calcarine sulcus.

At the whole brain level (Figure 7a), both *Stimulation paradigm* (*F*(1, 11) = 4.83, *p* = 0.05, *η*^2^ =0.31) and *Analysis procedure* (*F*(1,11) = 99.2, *p* < 0.001, *η*^2^ = 0.90) contributed to peak face-selective activity, without significant interaction. We found similar patterns within FG (Figure 7b) and ATL (Figure 7c). There were significant main effects of *Stimulation paradigm* (FG: *F*(1, 11) = 5.53, *p* = 0.038, *η*^2^ = 0.33; ATL, *F*(1, 11) = 4.01, *p*=0.07, *η*^2^ = 0.27) and *Analysis procedure* (FG: *F*(1,11) = 67.6, *p* < 0.001, *η*^2^ = 0.86; ATL: *F*(1,11) = 11.4, *p* = 0.002, *η*^2^ = 0.58) without any interaction. Hence, the results suggest that both factors contribute to the highest face-selective responses observed in key regions.

**Figure 7.**
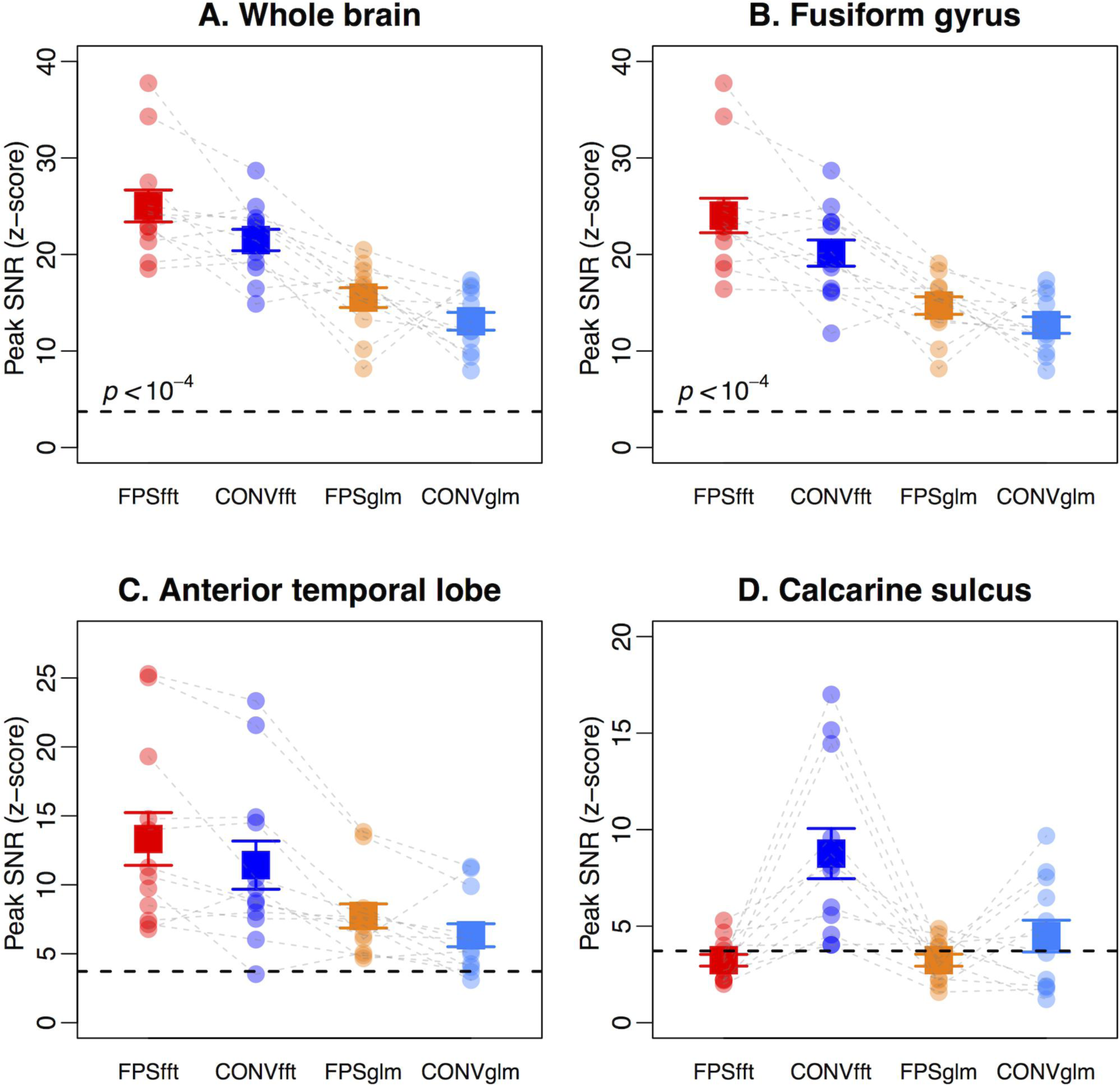
Respective contribution of analysis procedure and stimulation paradigm on peak face-selective response. (A) Peak face-selective response in the whole cortex. (B) Peak face-selective response in the fusiform gyrus; (C) Peak face-selective response in the anterior temporal lobe (D) Peak face-selective response in the Calcarine sulcus. *Note: FPSfft: FPS-face condition with FFT-based data analysis; CONVfft: CONV-face condition with FFT-based data analysis; FPSglm: FPS-face condition with GLM-based analysis; CONVglm: CONV-face condition with GLM based analysis. Each dot represents the value of one participant. The black dashed lines represent a threshold level of p < 10*^*-4*^, *uncorrected. The gray dashed lines connect the values from the same participant in different conditions. The solid squares represent the group means ± SEM. We combined homologous ROIs across hemispheres for this analysis.*

In the Calcarine sulcus (Figure 7d), the main effects of the *Stimulation paradigm* (*F*(1,11) = 11.1, *p* = 0.007, *η*^2^ = 0.50) and *Analysis procedure* (*F*(1,11) = 17.8, *p* = 0.001, *η*^2^ = 0.62) were modulated by a significant interaction (*F*(1,11) = 16.8, *p* = 0.002, *η*^2^ = 0.60). The pattern of results show that in the FPS-face condition, activity in the Calcarine sulcus was low and analysis procedure did not make any difference (*t*(11) = -0.02, *p* = 0.98). However, the FFT analysis was more sensitive than the GLM analysis in measuring “face-selective” activity in the Calcarine sulcus in the CONV-face condition (*t*(11) = 4.24, *p* = 0.001). The results indicate that the removal of low-level visual confounds is due to the stimulation parameters of the FPS-fMRI paradigm.

## Discussion

Many neuroscientists seek to understand brain function by characterizing and comparing neural responses within different cortical areas. Understanding what processes and representations occur in different brain areas requires well-justified and reproducible methods for defining these areas. In this context, functional localizers in fMRI have become invaluable tools, the most widely used comparing faces – arguably the most significant signal for human ecology – to non-face visual objects (Kanwisher et al., 1997; Kanwisher, 2017). This face localizer is critical to define regions in the human adult brain to explore further functions such as facial identity recognition or facial emotion decoding, but also to explore the neurodevelopment of this function (de Heering & Rossion, 2015; Gomez et al., 2017; Scherf, Behrmann, Humphreys, & Luna, 2007) and its impairment following sudden onset or long-life brain dysfunction (e.g., Avidan et al., 2014; Busigny et al., 2014;Susilo & Buchaine, 2013). However, it is fair to say that even after 20 years of use and development of functional fMRI localizers, severe limitations in sensitivity, objectivity, and reliability remain, explaining a large part of the great variability across studies in this field of research. Moreover, there is generally a trade-off in fMRI face localizers between the tight control of systematic low-level stimulus confounds, in which stimuli are equalized for low-level visual cues and a single or a few category of interest only are compared to faces (e.g., Gauthier et al., 2000; Rossion et al., 2012) at the expense of sensitivity and generalizability, and the use of more natural stimuli at the expense of specificity (e.g., Fox et al., 2009). Our FPS-fMRI approach, partly inspired from frequency tagging in electroencephalography, goes a long way towards resolving these issues.

Regarding sensitivity, our FPS-fMRI face localizer achieves much (i.e., twice) higher SNR in detecting face-selective neural responses overall than a conventional face localizer matched for duration and number of face stimuli presented. This advantage was found in every individual brain tested in the study. The increased SNR is due to both the stimulation paradigm and the model-free FFT analysis (Figure 7). The stimulation rate is optimized to give brain regions just enough time to selectively process each stimulus and to record a full category-selective response before the next stimulus from the same category appears (Retter & Rossion, 2016). It also maximizes the direct contrast between the category of interest, faces, and other stimuli, each face being forward- and backward-masked by non-face objects in the continuous visual stimulation stream. Due to the long run duration, frequency resolution of the fMRI signal spectrum following FFT is extremely high (0.0025 Hz). Hence, while the broadband noise spreads across numerous frequency bins, all the signal of interest falls in a tiny frequency bin (0.111 Hz), providing extremely high SNR (Regan, 1989). Increased sensitivity allows reducing acquisition time by a factor of two (see Supplemental Fig. S7), providing much more room for further exploration of specific functions of these regions, and to disclose all or most relevant face-selective regions in individual brains. The significant clusters in the ATL in the vast majority of individual brains with FPS-fMRI, despite signal dropout making it difficult to localize face-selective responses with conventional fMRI paradigms, illustrates this point.

The conventional way of estimating the BOLD response amplitude relies on the same HRF model being used for different regions of the brain and for different individuals. However, since there is a substantial amount of variation of BOLD hemodynamic response across brain regions as well as across participants, a uniform HRF inevitably leads to erroneous estimates (Handwerker, Ollinger, & D’esposito, 2004). This matter is even worse for modeling hemodynamic response in higher-level regions, since commonly used HRF models are usually validated from measurements of specific regions and stimulation, such as the primary visual cortex to simple visual stimuli (e.g., Boynton et al., 1996) or the primary motor cortex to finger-tapping (Boxton et al., 1998). In contrast, the model-free FFT analysis afforded by periodicity in the FPS-fMRI design leads to a more objective definition and quantification of a response of interest (i.e., amplitude exactly at the pre-defined frequency), but also a fair comparison of this response across different brain regions and participants. Note that a key difference with previous frequency-based fMRI analyses (Koenig-Robert, VanRullen & Tsuchiya, 2015; McCarthy et al., 1995; Puce et al., 1995; 1996; Wang et al., 2014, 2015) is that category-selective neural responses are directly reflected as amplitude of the stimulation frequency here, without post-hoc subtraction procedure.

Importantly, our approach also achieves high test-retest reliability in localizing face-selective areas. Although reliability is the pedestal of any scientific research, the results of fMRI research are generally less reliable than many researchers implicitly believe (Bennett & Miller, 2010). Here, across the whole brain or within functionally defined face-selective regions, we achieve an average of 80%-90% overlap of spatial activation across runs, which is to our knowledge the highest test-retest reliability observed in this field of research, and beyond. This high reliability makes the current paradigm desirable not only for fundamental scientific research, but, importantly, for clinical purposes, such as presurgical functional localization. This reliability is more impressive considering that the specific exemplar images are selected randomly for each presentation, and thus not presented in the same order, or repeated exactly the same number of times in each run. Precisely, we attribute the high reliability of the approach to the measure of a process, perceptual categorization, which is repeatedly sampled across a wide variety of stimuli and contexts (i.e., non-face stimuli) within each run, increasing invariance, i.e., robustness, of the response of interest.

Critically, the increase in sensitivity afforded by the current paradigm does not come at the expense of a decrease of specificity, which is often the case when natural stimuli are uncontrolled for low-level properties (Fox et al., 2009) or in block designs with one-back tasks differing in difficulty and attentional resources between face and object images (see discussion in Rossion et al., 2012). In fact, overall, there are no more voxels activated in our paradigm than in the conventional paradigm with the same stimuli: the significant activation increase in higher-level regions such as the ATL, defined as face-selective in previous fMRI studies of face-selectivity and human intracranial recordings (Rajimehr et al., 2009; Nasr & Tootell, 2012; Puce et al., 1999; Jonas et al., 2016) and considered as a key player in face perception, is mainly compensated by the lack of “face-selective” responses in the primary visual cortex with the FPS approach compared to the conventional localizer. Note that the same stimuli are used in both designs and FFT analysis *per se* does not account for the removal of low-level visual cortex activation. Hence, removal of low-level visual confounds is rather due to the fast periodic stimulation paradigm. In a conventional block design, a mere minority – say 10% – of images of the same category with specific low-level visual attributes (e.g., increased power in low spatial frequencies) could well lead to average differences between (face and object) stimulation blocks. Moreover, with natural images as used here, populations of neurons responding to living things in general, for instance, would contribute to “face-selectivity” in a block design (or in studies comparing faces to houses or cars only, e.g., Loffler et al., 2005; Rossion et al., 2012). However, the periodicity constraint in FPS-fMRI eliminates such confounds (Rossion et al., 2015) because each face image contrasts with different object images throughout the sequence, and the face bursts are made of a few (different) exemplars only. Hence, a subset of identical biased image contrasts would have to be systematically present in each face burst to lead to a periodic response due to such biases. A major strength of the present approach is the ability to preserve naturalness of the images while removing low-level visual contributions, as demonstrated here by the absence of face-selective response when these images are phase-scrambled.

The principles of FPS-fMRI validated here with a functional face localizer could be applied straightforwardly to refine our knowledge of the functions of these regions with respect to face identity and facial expression processing for instance (Haxby et al., 2000; Duchaine & Yovel, 2015), to map cortical areas selectively coding for visual linguistic material (e.g., letter strings and words, Lochy et al., 2015) in the VOTC (Wandell, 2011), or to other modalities such as auditory stimulation (Zatorre et al., 2002). More generally, our approach is able to objectively track fast processes occurring in a dynamic visual stream even with a slow temporal resolution method such as fMRI. This is important for providing direct comparisons with neural signals obtained in similar paradigms used with other imaging modalities, such as (intracranial) electroencephalography (Jonas et al., 2016), and also open new possibilities for defining functions with other low temporal resolution methods such as calcium imaging (Grienberger & Konnerth, 2013) in human and nonhuman brains.

## Author contributions

X.G. and B.R. designed the research, X.G. and F.G. conducted the research, X.G. analysed the data, X.G., F.G., and B.R. wrote the manuscript.

The authors declare no conflict of interest.

## Acknowledgements

This work was supported by European Research Council Grant facessvep 284025 to B.R., and an UCL/Marie-Curie postdoctoral fellowship to X.G. We thank Valérie Goffaux, Corentin Jacques, Jacques Jonas, Kirsten Petras, and Talia Retter for their invaluable comments on an earlier version of this paper.

## Methods

### 1. Participants

Twelve adults (8 females, mean age = 30.1 ± 5.4 years, age range = 24 to 42 years) from the York University (Canada) community participated in the fMRI experiment. All of the participants had normal or corrected-to-normal vision, and were right-handed (Oldfield, 1971). None of the participants reported any history of psychiatric or neurological disorders, or current use of any psychoactive medications. The study was approved by York University Research Ethics Board (certificate #: e2014-155). We obtained informed written consent from all the participants prior to the experimental sessions and they received $50 for their participation in the study.

### 2. Stimuli

The stimuli consisted of 100 face images and 200 non-face images (e.g., Figure 8, the whole set of face and non-face images can be obtained upon request). The face images were digital photographs of 100 different individuals who were non-famous relatives, friends and colleagues of the researchers of the *Face Categorization Lab* of the University of Louvain, Belgium. Therefore, they were unfamiliar to the participants. Each photograph contained one human face. The photographs were originally taken for personal purposes, and were given to the researchers with a completed consent form to use these photographs for research purposes and display them. They contain a natural range of variation in size, pose, and expression of the faces depicted in the photographs and in lighting and background. The non-face images consist of 200 photographs of scenes, objects, and animals. As in the face images, the non-face images also contain a natural range of variation in the composition and lighting of the images. The face images have a mean grayscale intensity value of 115.0 ± 1 and a mean contrast value of 0.49 ± 0.11. The non-face object images have a mean grayscale intensity value of 115.2 ± 0.9 and a mean contrast value of 0.46 ± 0.12. On average, there is no statistical difference between the two sets of images on either the grayscale intensity value or the image contrast. Pixel-wise averaging of either all the face images or all the non-face images did not reveal any identifiable structure (Figure 8, the right most column). We created a Fourier phase scrambled version of each image (e.g., Figure 8, bottom row) by randomizing the phase of the original images (Sadr & Sinha, 2004), as used in previous EEG studies (e.g., Rossion et al., 2015). At a global level, these images contain the same low-level visual information (i.e. power spectrum) of the original images, but without any recognizable structure.

**Figure 8.**
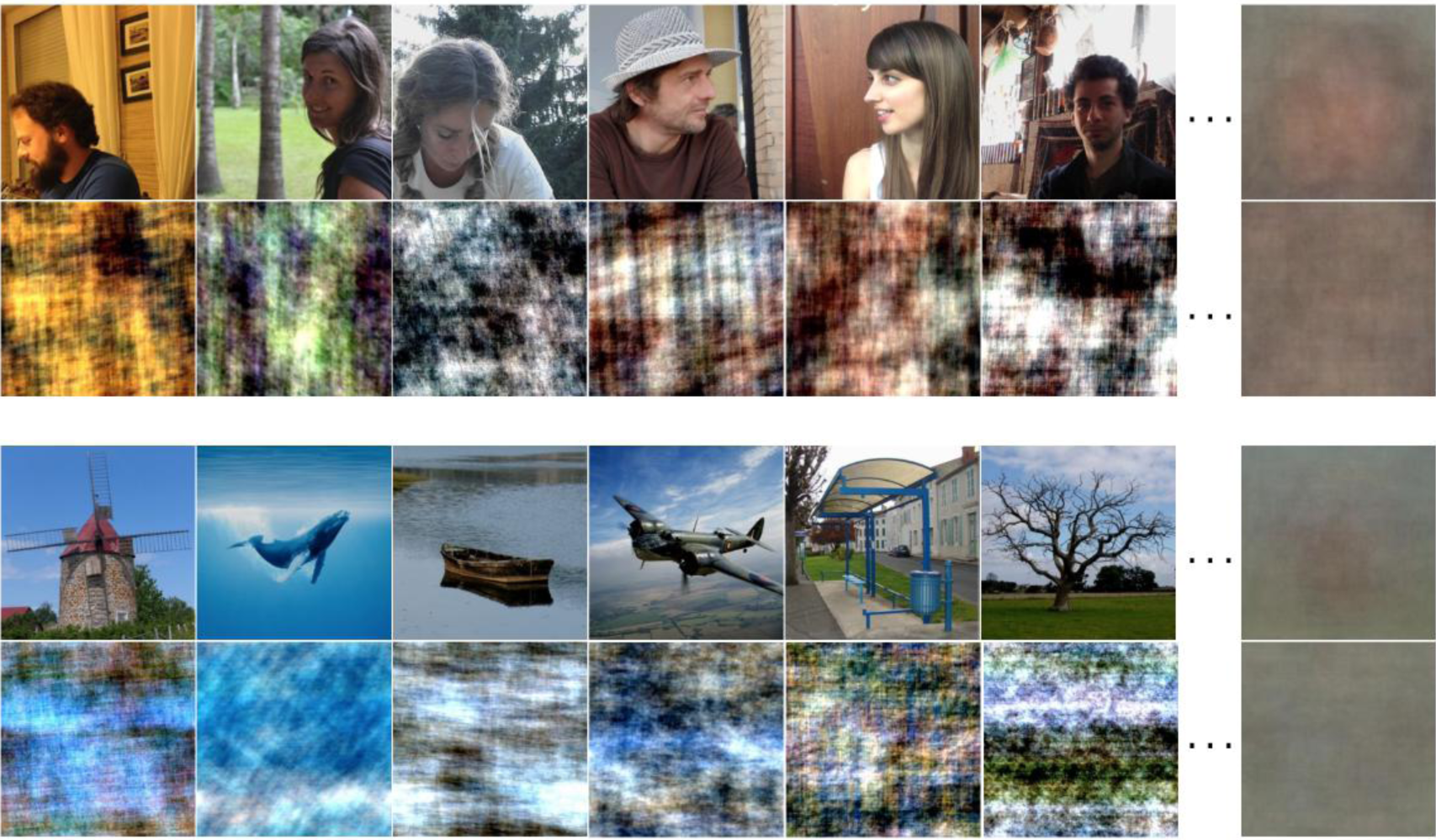
Example stimuli. Both faces and objects contain a wide range of variation in the composition, color, and lighting. Such high variability reduces the contribution of specific low-level visual cues to perceptual categorization while preserving naturalness of the images (Rossion et al., 2015). No amplitude spectrum equalizing was performed on these images. Fourier phase-scrambled versions of the images (below each face and object image) were also created. The right-most column contains pixel-wise averaged images of all the faces, all the objects, and all the phase scrambled images within each category. No recognizable structure is seen in the average images.

### 3. Stimulation procedure

The images were back-projected in full color onto a projection screen by an MRI compatible LCD projector and viewed by the participant through a mirror placed within the RF head coil at a viewing distance of 43 cm. They extended 14.6×14.6 degree of visual angle at the viewing distance (or 11×11 cm on the screen). The remaining area of the screen was set to a uniform gray background. The whole experiment procedure was controlled through a stimulation program running in Java, which also collected behavioral responses.

#### 3.1 Main stimulation: FPS-fMRI

As shown in Figure1a, in the FPS sequence, the images were displayed at a base rate of 6 Hz (i.e., 6 natural images/sec, the blue bins in the figure), thus with a stimulus onset asynchrony (SOA) of 166.7 ms (10 screen refresh cycles at a refresh rate of 60 Hz), allowing one gaze fixation per image. Images were contrast modulated by a sinusoidal function so that each image appeared at 0% contrast, reached 100% contrast at the 6^th^ frame and then dropped its contrast to 9.55% at the 10^th^ frame (Figure1b). The same sinusoidal contrast modulation has been used in previous EEG studies (e.g., Rossion & Boremanse, 2011), in particular with this paradigm (e.g., Rossion et al., 2015). Although virtually identical amplitude results can be observed in EEG when stimulating with 50% duty-cycle squarewaves and sinewaves (e.g., Retter & Rossion, 2016), using sinewaves has several advantages. First, since images at low contrast are visible, the visual stimulation is almost always present and gives the observer a continuous perceptual stimulation; second, it is smoother as a visual stimulation than abrupt changes (i.e., squarewaves).

Every 9 seconds, a set of 7 faces (referred to as a *face burst*, covering 2.167 secs, the red bins in Figure 1a) appeared at a rate of 3 Hz, i.e. alternating with non-face images. We alternated between faces and objects repeatedly during a face “burst” for three reasons. First, alternation is preferred to a block of identical events to reduce category-specific adaptation (e.g., Kovacs et al., 2006) and to ensure that the temporal distance between two faces (here 333 ms) captures the bulk of the face-selective response in the human brain as estimated with scalp EEG (Retter & Rossion, 2016). Second, a repeated alteration should strengthen the differential neural response between faces and objects. Third, we reduced the likelihood that a participant misses the alteration, which may have happened if there was only a single alteration every 9 seconds. Each run had a length of 396 seconds so that the face burst appeared 44 times at a fixed frequency of 1/9Hz (i.e., 0.111 Hz, referred to as the *face stimulation frequency*). For each presentation, an image was drawn from the corresponding image set (face or non-face) according to a random order. When all the images in the respective sets had been presented, a new random order was generated and the images were drawn according to this new random order. So across all the runs and all the subjects, the order of image presentation were always different as it was randomly generated online. In total, in one FPS-fMRI run, face images appeared 7×44 = 308 times while non-face images appeared 2,068 times.

#### 3.2 Phase-scrambled image stimulation

The design included a scrambled image condition, which had exactly the same design as the FPS-face sequence but with phase-scrambled face and non-face images. We refer this condition to as FPS-scrambled.

#### 3.3 Conventional face localizer

To test the sensitivity and selectivity of the FPS-fMRI approach, we included a conventional fMRI block design functional localizer (“face localizer”, e.g., Kanwisher et al., 1997), and referred it to as CONV-face condition. In this design, faces and non-face images appeared in alternating 18-sec blocks without resting in between. Such a design is optimal for estimating the contrast between two conditions (Smith et al., 2007; Maus et al., 2010). We matched the conventional face localizer to the FPS-face runs for two critical aspects: run duration and total number of face images presented. Specifically, the run length is exactly the same as in the FPS-face sequence (396 secs) with 11 repetitions of the face and non-face blocks. In total, face images appeared for 28×11 = 308 times, which matched the number of the appearance of faces in the FPS-face runs. However, in the conventional face localizer, the non-face images also appeared for 308 times, as in typical block design studies. Within each 18 sec block there were 28 images displayed at 1.56 Hz. Each image appeared for 643 msec with its contrast ramped up from 80% to 100% and then dropped back to 80% following a sinusoidal function. This way, images were presented successively, without a blank interval. The modulation of the contrast provided a relatively smooth transition between images (Supplemental movie S2 for an example).

### Behavioral task and order of conditions

In all the three conditions, the participants performed the same behavioral task, orthogonal to the measure of interest. They were instructed to press a predefined key on an MRI compatible response pad using the right index finger when they detected color changes of the central crosshairs superimposed on the images (Rossion et al., 2015). The crosshairs extended a visual angle of 1.2 degrees in the center of the screen. During each run, the color of the crosshairs changed from black to white for 200 ms, for a total of 70 times with the interval between two changes randomized while keeping above a minimal interval of 2 secs. All participants achieved high accuracy (mean accuracy across conditions range from 0.927 to 0.993) in the behavioral task with no significant difference among the three conditions in either accuracy (*F*2,11 = 3.53, *p* = 0.07, Greehhouse-Geisser corrected) or correct response time (*F*2,11 = 1.55, *p* = 0.23).

Each participant started the first session with one run of each condition (FPS-face, FPS-scrambled, CONV-face) in a random order. A high-resolution anatomic image was obtained after the first session. After the anatomic image, each participant continued with the second session of one run of each of the three conditions in a pseudo-random order with the first run in the second session being a different condition from the last run in the first session. The participants took a short break after finishing the second session. After the break, they continued with the third session with only one run of the FPS-face condition and one run of the CONV-face condition in a pseudo-random order, with the first run in the third session being a different condition from the last run in the second session. In total, each participant had three FPS-face runs, three CONV-face runs, two FPS-scramble runs, and one anatomic run. The whole experiment took about one hour and thirty minutes.

### MR Image acquisition

We acquired the MRI images using a 3T Siemens Magnetom Trio system (Siemens Medical System, Erlangen, Germany) with a 32-channel head coil. Anatomic images were collected using a high-resolution T1-weighted magnetization-prepared gradient-echo image (MP-RAGE) sequence (192 sagittal slices, TR = 2,300 ms, TE = 2.62 ms, voxel size = 1 mm isotropic, FA = 9°, FoV = 256 × 256 mm^2^, matrix size = 256 × 256, parallel scanning mode = GRAPPA, accelerate factor =2). The acquisition time for the anatomic scan was 321 seconds. Functional images were collected with a *T*_*2*_*\** weighted gradient-echo echoplanar imaging (EPI) sequence (TR = 1,500 ms, TE = 30 ms, FA = 62°, voxel size = 3 mm isotropic, FoV = 192 × 192 mm^2^, matrix size = 64 × 64, interleaved, parallel scanning mode = GRAPPA, accelerate factor = 2), which acquired 25 oblique-axial slices covering the whole occipital lobe and the whole temporal lobe. The acquisition time for each functional run was 414 seconds.

### 6. MRI/fMRI Analysis

#### 6.1 Preprocessing

The functional runs were motion-corrected in reference to the average image of the first functional run of the experiment using a 6-degree rigid body translation and rotation via an intra-modal volume linear registration using the FMRIB Software Library (FSL, version 5.0.8, Smith et al., 2004). The motion-corrected images were spatially smoothed with a Gaussian kernel with a moderate size (3 mm FWHM) to increase overlap of regions across runs and reduce noise level while keeping a high spatial resolution (Worsley et al., 1996).

#### 6.2 Amplitude of category-selective response

For each run with the FPS-fMRI paradigm (FPS-face and FPS-scrambled), we removed linear trends from the preprocessed time series data of each voxel and converted the time series data to percentage of BOLD signal change by dividing the time series of each voxel by its mean signal intensity (Figure 2b). We then performed Fast Fourier Transform (FFT) to obtain the amplitude spectrum (Figure 2c) of the time series. To gauge the strength of the BOLD response at the face stimulation frequency (the signal) relative to the noise, we converted the amplitude of the face stimulation frequency (0.111 Hz) to a z-score as in previous studies (McCarthy et al., 1994; Puce et al., 1995). Intrinsically, a z-score is a measure of SNR. Therefore, we referred the z-scores to as SNR of the face-selective neural responses.

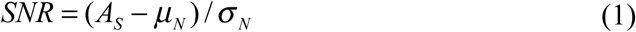

where *A*_*S*_ is the amplitude of the face stimulation frequency, *μ*_*N*_ is the mean and *σ* _*N*_ is the standard deviation of the amplitude of 40 neighboring frequencies (20 on each side, e.g., Rossion et al., 2015, Jonas et al., 2016; see methods, Figure 2d). This procedure is applied to each voxel independently.

#### 6.3 Direction of category-selective response

A voxel having a high SNR value at the face stimulation frequency responded differentially to faces and objects, but not necessarily more to faces than objects. Indeed, it is possible for a voxel to achieve a high SNR at the face stimulation frequency if the BOLD response systematically *decreases* (i.e., deactivates) to the appearance of faces relative to the objects. To define the direction of category-selective responses in the FPS-face condition for each voxel, we calculated the averaged BOLD response time course across all the cycles (44) during the face stimulation run. We then calculated the differences at time points two to six relative to time point one within the averaged time course. We used the sign of the summed differences as the direction of category-selective response. A positive value means activation while a negative value means deactivation to the presence of faces. We created a sign map and applied this map to a thresholded SNR (z-score) map (Figure 9). We obtained the final category-selective response map contains only voxels that have increased BOLD response to the presence of faces.

**Figure 9.**
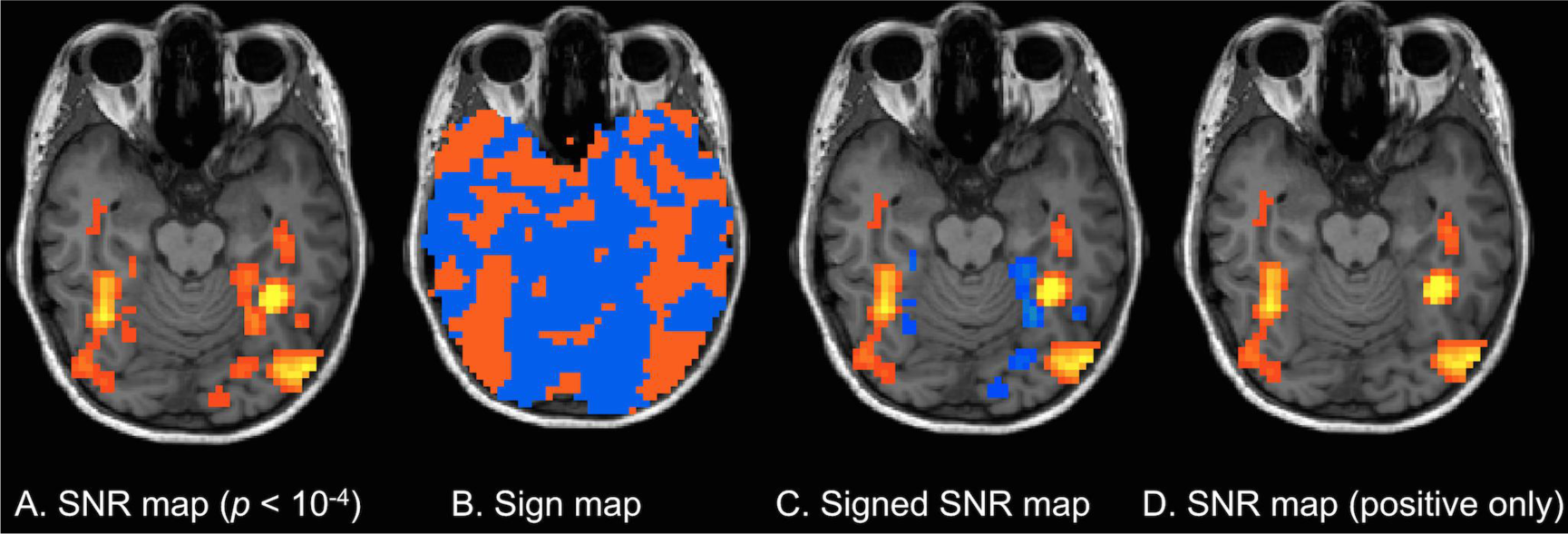
Sign masking of the FPS-face runs (in one example brain) (A) Unsigned SNR map with voxels that are above threshold of *p* < 10^-4^, uncorrected. (B) Sign map calculated from averaged BOLD response time course across all (44) cycles, orange for positive sign (activation) and blue for negative sign (deactivation). (C) A signed SNR map created by applying the sign map in (B) to the map in (A). (D) SNR map with negative values removed.

#### 6.4 Conventional face localizer

For each CONV-face run, following a standard procedure (e.g., Weiner & Grill-Spector, 2010), we estimated the voxel-wise BOLD response amplitudes to face blocks and non-face blocks by fitting the preprocessed time series data with a general linear model (GLM), convolved with a hemodynamic response function (canonical HRF, SPM8) with temporal derivatives, autocorrelation model type AR(1), and with nuisance regressors including 6 movement parameters. We conducted a linear contrast to obtain the *t* statistical map where the BOLD responses to faces were greater than to non-face images. By nature, the *t* values are a measure of *SNR*, since the *t* values are calculated by dividing the difference of activation amplitudes between faces and non-face images (signal) by the estimated standard error of the activation (noise). To make the SNR directly comparable between conditions, we converted the *t* values to z scores by calculating the corresponding *p* values from the *t* distributions.

#### 6.5 Using data from all runs

In the current experiment, we collected data from 3 runs in the FPS-face condition, from 3 runs in the CONV-face condition, and from 2 runs form the FPS-scrambled condition. By using the data from all the runs within each condition, we increased statistical power. For the FPS-face condition, across runs, the responses to the periodic face stimulations in a given population of neurons have the same phase, while any noise from a periodic source (e.g., pulse, breathing) could have different phases across runs. Therefore, we averaged the time series across the three FPS-face runs to increase the signal-to-noise ratio, similarly to the use of this approach in electrophysiology (Regan, 1989). Since the same principal applies to the FPS-scramble condition, we averaged time series across the two FPS-scramble runs. For the conventional localizer, we run the same GLM analysis as with individual runs, but with the time series from all three runs. In this case, run is added to the GLM analysis as a fixed effect. By using data from all three runs in the GLM analysis, we increased the degree of freedom in the Face>Non-face contrast from 241 to 723. Therefore, we increased power to detect any differences between neural responses to faces and non-face objects. With the same procedure as with individual runs, we created a z-map for each condition with data from all runs for each participant.

#### 6.6 Group analysis

We used the individual z-maps derived from data based on all runs for each condition to calculate an average group z-map for each condition. We used cortical surface-based averaging algorithms in FreeSurfer, which has been show to yield a better alignment across subjects than volume-based normalization methods (Fischl et al., 1999). The surface-based approach morphs each participant’s cortical surface reconstructed from high-resolution anatomical scan to an averaged spherical surface of all the 12 participants using a best-fit sulci alignment. Individual z-maps were then interpolated onto the average sphere and averaged across participants. The averaged z-scores are no longer from a standard normal distribution. Instead, they have a standard deviation of 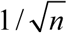 (*n* = 12, number of participants). Therefore, we adjusted the z threshold with the same factor 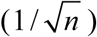. Because group averaging can be affected by extreme values, we applied a conservative threshold (*p* < 10^-6^, uncorrected, equivalent to a Bonferroni corrected *p* < 0.05) to identify areas that are face-selective at a group level.

#### 6.7 Region of interest (ROI) based individual analysis

Previous studies have shown a considerable amount of variability across individuals in both brain structure and functional localization (Frost and Goebel, 2012; Zilles and Amunts, 2013), particularly in face localizers (Rossion et al., 2012; Zhen, et al., 2015). Therefore, it is critical to also analyse the face-selective responses at an individual level. We first created an average cortical surface and mapped the individual face-selective activity to the average cortical surface as in the group analysis. We then created 20 anatomically defined ROIs (10 in each hemisphere) on the average cortical surface. The ROIs were selected based on the results of the group analysis and were defined by an automatical parcellateion scheme by Freesurfer. The parcellation algorithms are based on anatomical rules and have good concordance with manual labels (Destrieux, Fischl, Dale, Halgren, 2010). The selected ROIs include: *inferior occipital gyrus, calcarine sulcus, cuneus, parieto-occipital sulcus, precuneus, fusiform gyrus, lingual gyrus, anterior occipito-temporal sulcus, superior temporal sulcus, and inferior frontal sulcus*. In addition, we manually drew on the cortical surface a ROI covering the anterior collateral sulcus. For the superior temporal sulcus, we further divided it into an anterior portion and a posterior portion with the boundary defined by the posterior tip of the hippocampus (Kim et al., 2000). We combined fusiform gyrus and lateral occipito-temporal sulcus into a single ROI, since the activation in the fusiform gyrus extends laterally into the sulcus. We also combined the anterior occipito-temporal sulcus and the anterior collateral sulcus into a single ROI, representing a face-selective area in the ATL. Figure 10 shows the ROIs on an inflated cortical surface.

**Figure 10.**
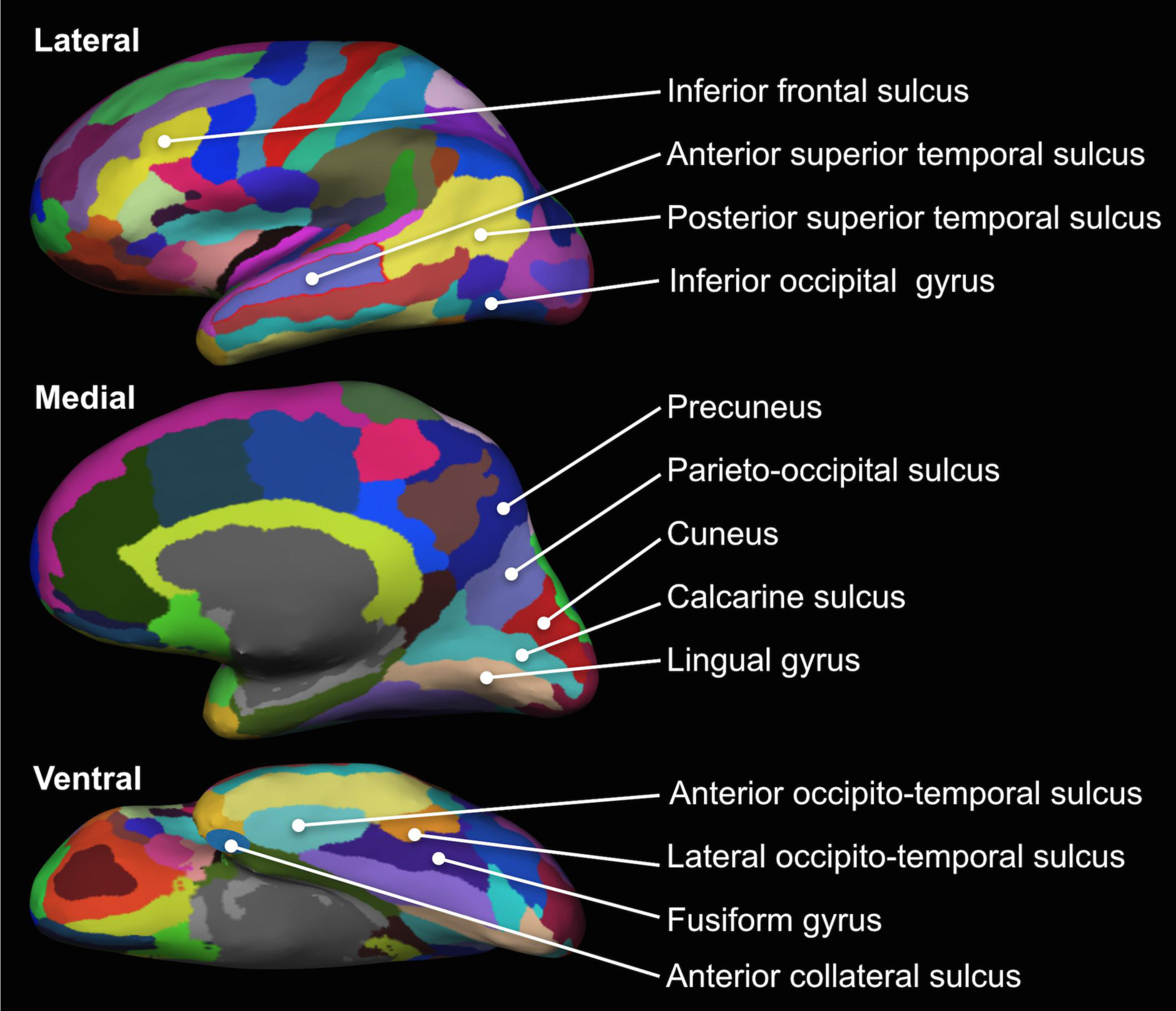
Anatomically defined regions of interests (ROIs) The ROIs were selected based on the results from the group analysis. The ROIs were automatically parcellated by Freesurfer based on reconstructed cortical surface constructed from high-resolution anatomic scans. We divided the superior temporal sulcus into an anterior portion and a posterior portion with the boundary defined by the posterior tip of the hippocampus. We combined fusiform gyrus and lateral occipito-temporal sulcus to be one ROI since the activation in the fusiform gyrus extends laterally into the sulcus. We manually labeled anterior collateral sulcus and combined it with anterior occipito-temporal sulcus to form the ATL ROI.

#### 6.8 Test-retest reliability

To quantify the overlap between face-selective voxels identified in different runs, we calculated a consistency score using the Dice coefficient:

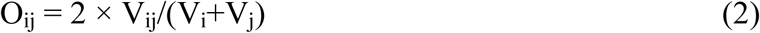

Where O_ij_ is the consistency score between run i and run j, V_ij_ is the number of super-threshold voxels in both runs, V_i_ is the number of super-threshold voxels in run i, and V_j_ is the number of super-threshold voxels in run j.

Within each condition, we calculated the consistency scores between run 1 and run 2, between run 1 and run 3, and between run 2 and run 3. We then averaged the three consistency scores to get a consistency score for that condition. For the analysis within functionally defined right FFA, since the data from the first run is used to define the ROI, we calculated the Dice coefficient with data from run 2 and run 3.

### Respective contribution of analysis procedure and stimulation paradigm

One advantage of the FPS paradigm is that it does not rely on the GLM framework. Instead, neural response amplitude is measured by applying FFT to the neural response time course. As explained earlier, the FFT approach has high SNR as the signal is only affected by narrow-band noise. To investigate the potential impact of the two data analysis approaches (GLM vs. FFT), we performed GLM analysis to the FPS-face data. Correspondingly, we performed FFT analysis to the data from the CONV-face runs.

#### 7.1 GLM analysis of the FPS-face data

As with the GLM analysis with the data from the CONV-face runs, we modeled BOLD response with two event types, faces and non-face objects, the two events alternate for 44 times within each scan run. Each face event lasts for 2.167 secs, while each non-face event lasts for 6.833 secs with no gap in between events. We included in the model data from all three FPS-face runs and modelled run as a fixed effect. A linear contrast (t-test) was constructed to compare the response amplitudes to faces and to non-face objects. The resulting *t* values were converted to z-scores for further comparison.

#### 7.2 FFT analysis of the CONV-face data

In the conventional localizer, we had 18-sec face blocks and 18-sec non-face object blocks alternating 11 times within each scanning run. Such a periodic presentation of the blocks allowed us to analyse the neural responses in the frequency domain. Specifically, both face-selective and non-face selective neural responses should have high amplitudes at 1/36 Hz. Similar to the FPS-face condition, we obtained signs for each voxel using the averaged BOLD response time course across all (11) cycles. Specifically, within the averaged time course, we calculated the difference between the sum of percent BOLD signal change in the first half cycle (first 12 time points) and the sum of percent BOLD signal change in the second half cycle (last 12 time points). Since face blocks were always presented before non-face blocks in all the cycles, a positive difference score suggests activation to faces while a negative difference score suggests deactivation to faces. We applied the signs to the threshold SNR map and only kept the voxels with positive values.

---

**Supplemental Fig. S1:**
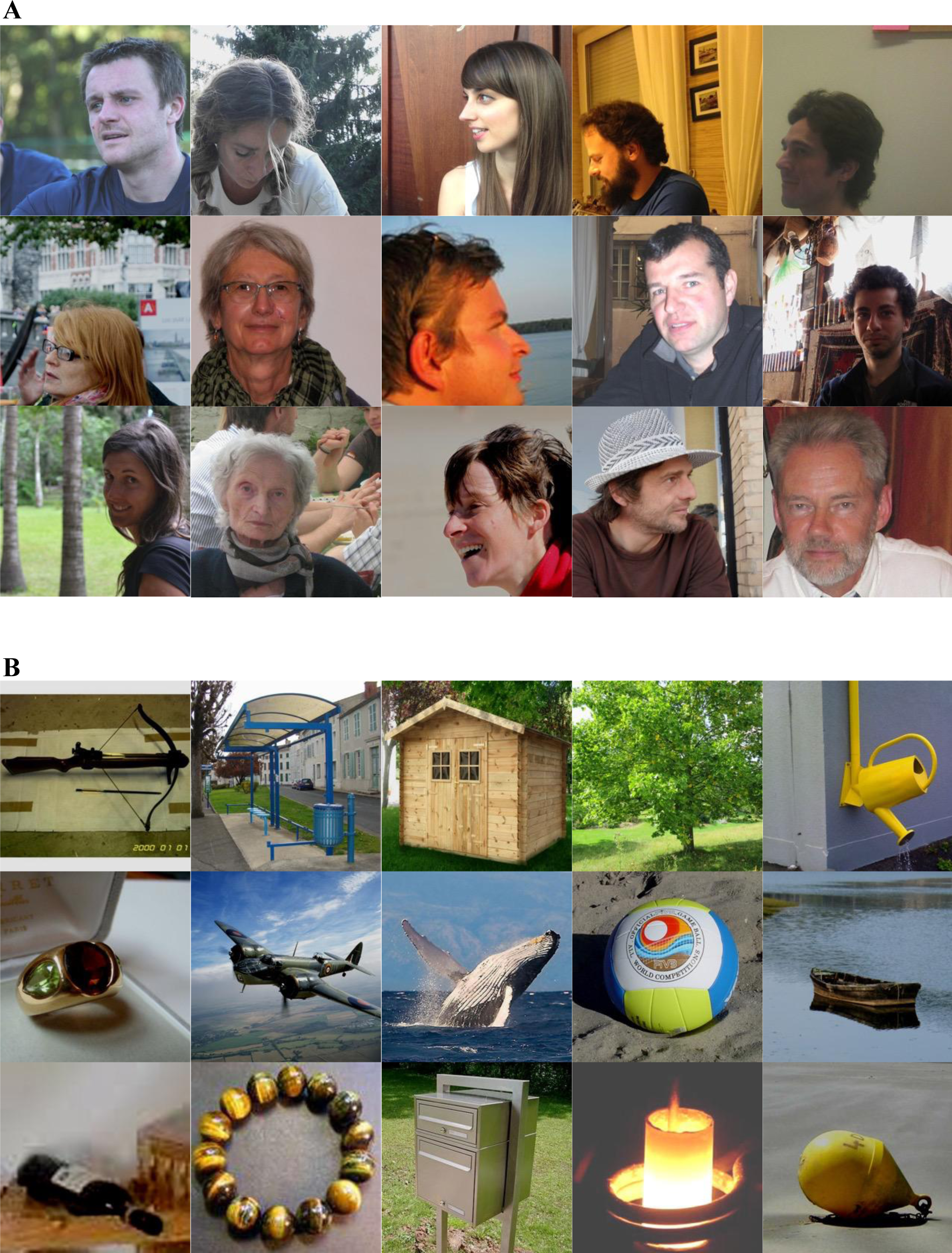
Examples of natural images. Related to Figure 1. Sets of face images (A) and non-face object images (B) without equalizing their low-level visual properties such as colour, luminance, and contrast. Both sets contain a natural range of variation.

**Supplemental Fig. S2:**
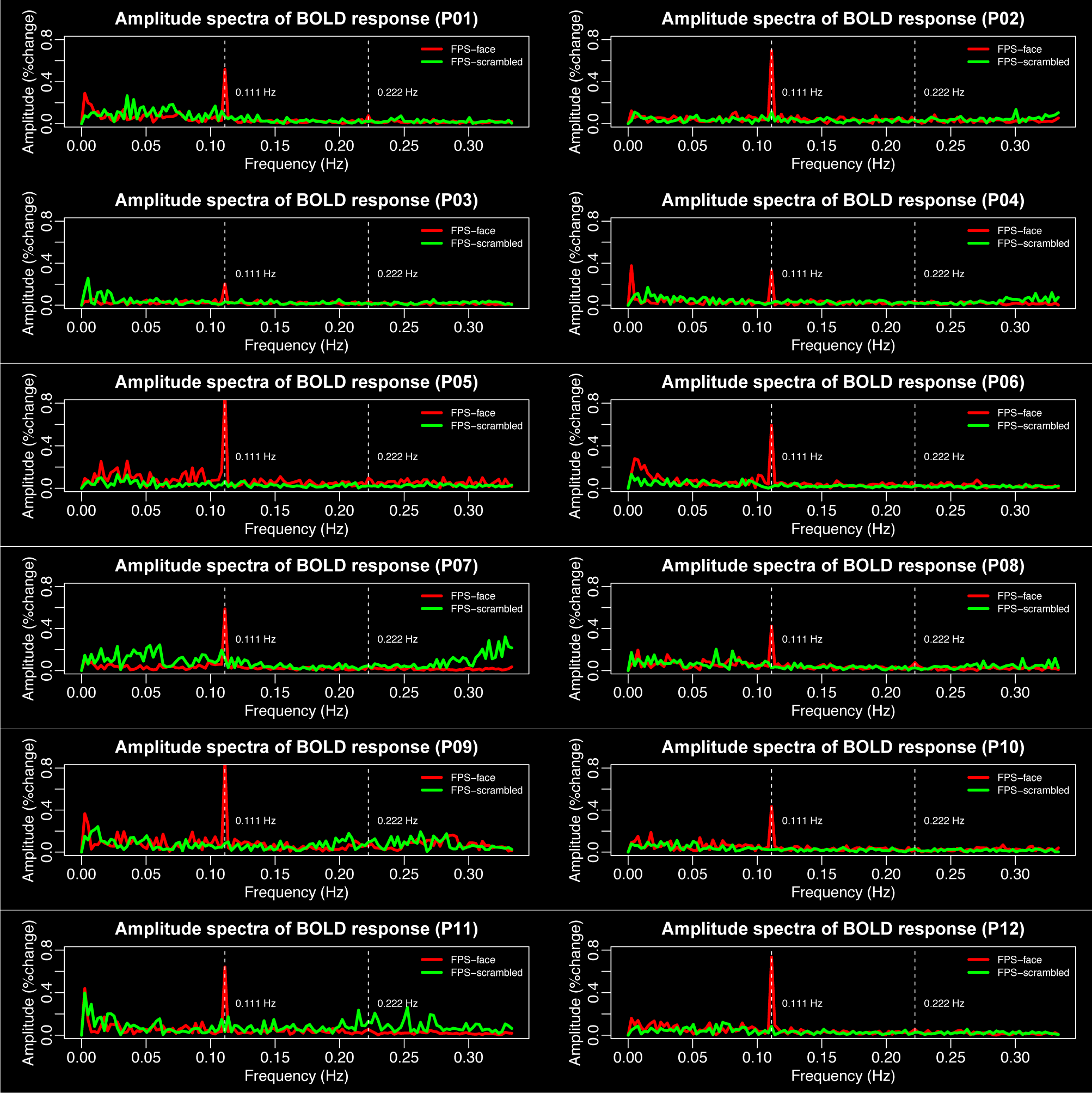
Amplitude spectra of BOLD response at the peak right FFA voxel of each participant (P01 to P12). Related to Figure 2.

**Supplemental Fig. S3:**
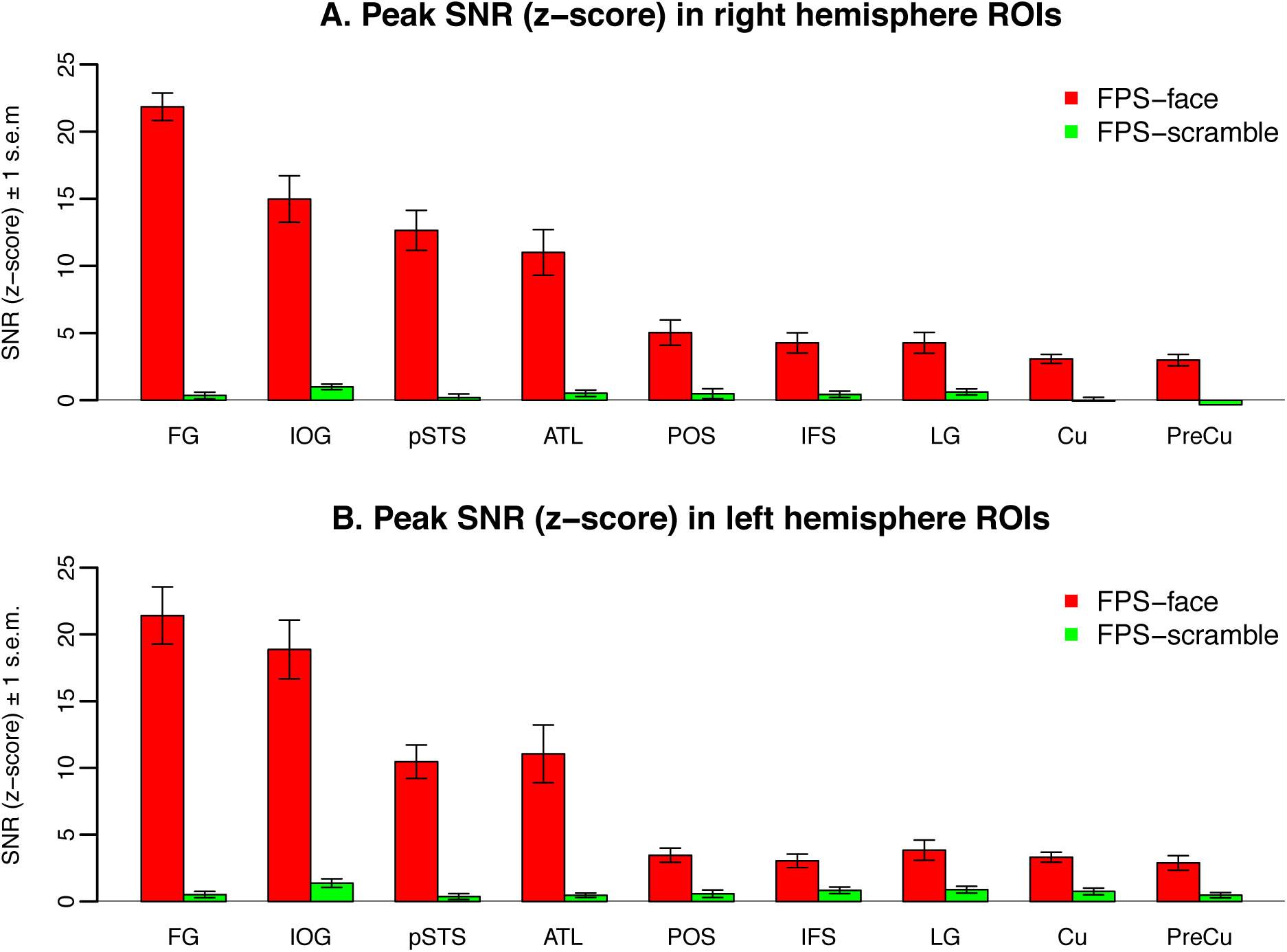
Signal-to-noise ratio (SNR) of the peak face-selective voxel in anatomically defined areas of interests (ROIs). Related to Figure 2. Signal-to-noise ratio (z-score) of the face stimulation frequency (0.111Hz) of the peak face-selective voxel identified by the FPS-face localizer in anatomically defined areas of interests in the right (A) and left (B) hemispheres. The bars represent the average value across 12 participants in the FPS-face condition (red) or the FPS-scrambled condition (green). Error bars represent 1 s.e.m. FG: fusiform gyrus; IOG: inferior occipital gyrus; pSTS: posterior superior temporal sulcus; ATL: anterior temporal lobe; POS: parietal-occiptal sulcus; IFS: inferior frontal sulcus; LG: lingual gyrus; Cu: cuneus; PreCu: precuneus.

**Supplemental Fig. S4:**
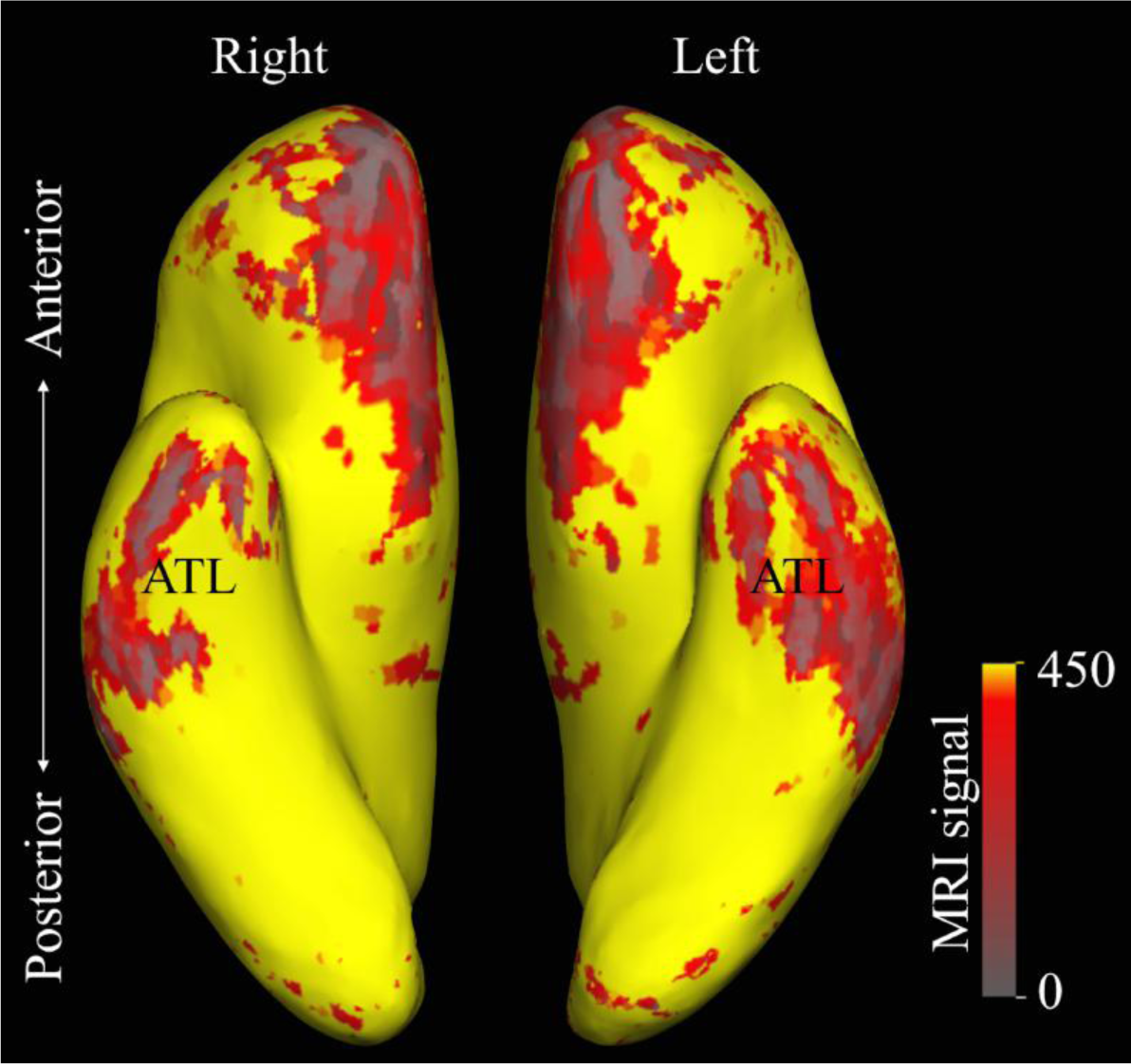
MRI signal drop-out in the anterior temporal lobe. Related to Figure 3. An example of MRI signal drop-out in the brain of one example participant. Weak MRI signal was recorded in the anterior temporal lobe (ATL) in both hemispheres as a result of magnetic susceptibility caused by the ear canals.

**Supplemental Fig. S5:**
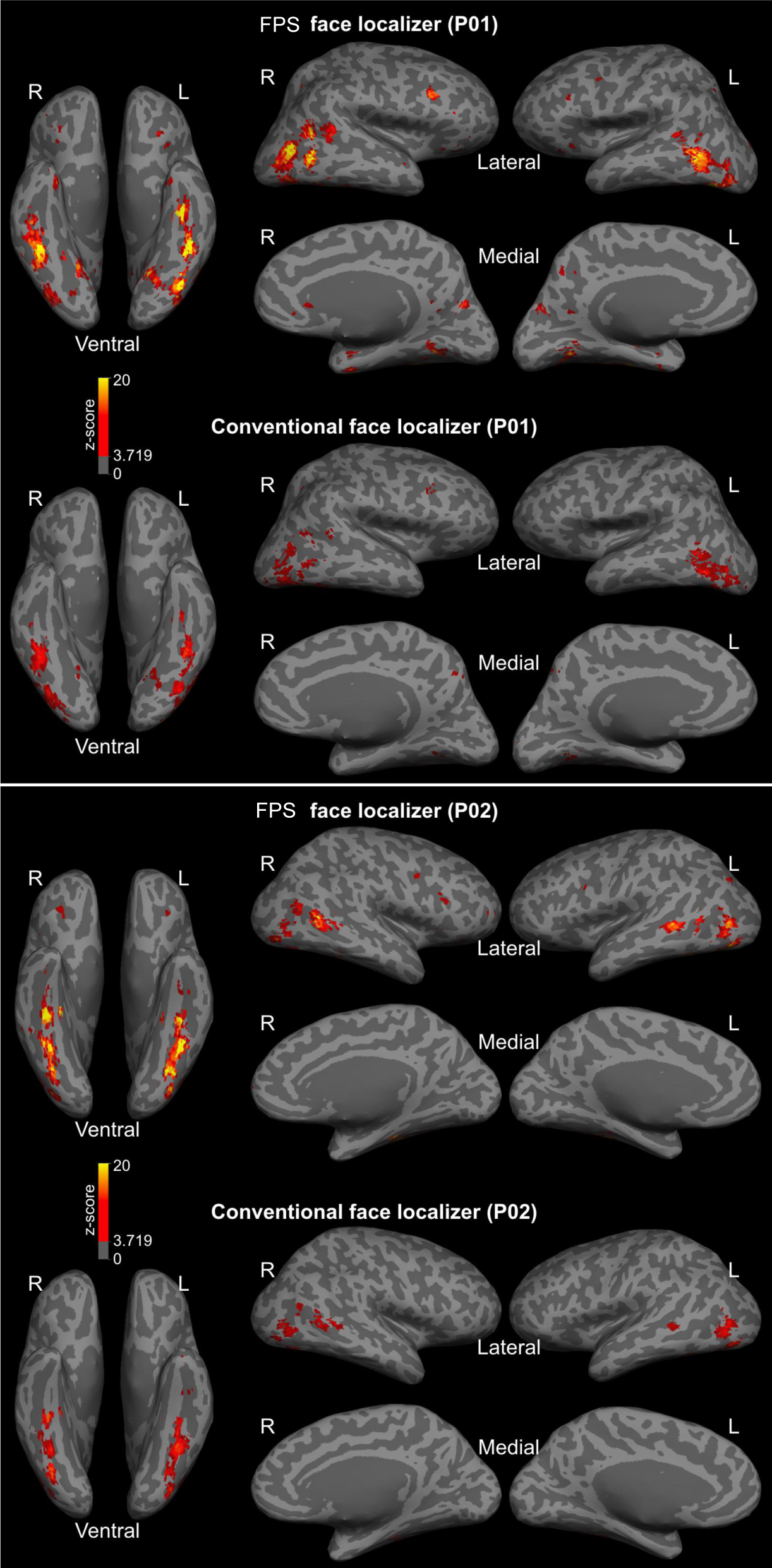
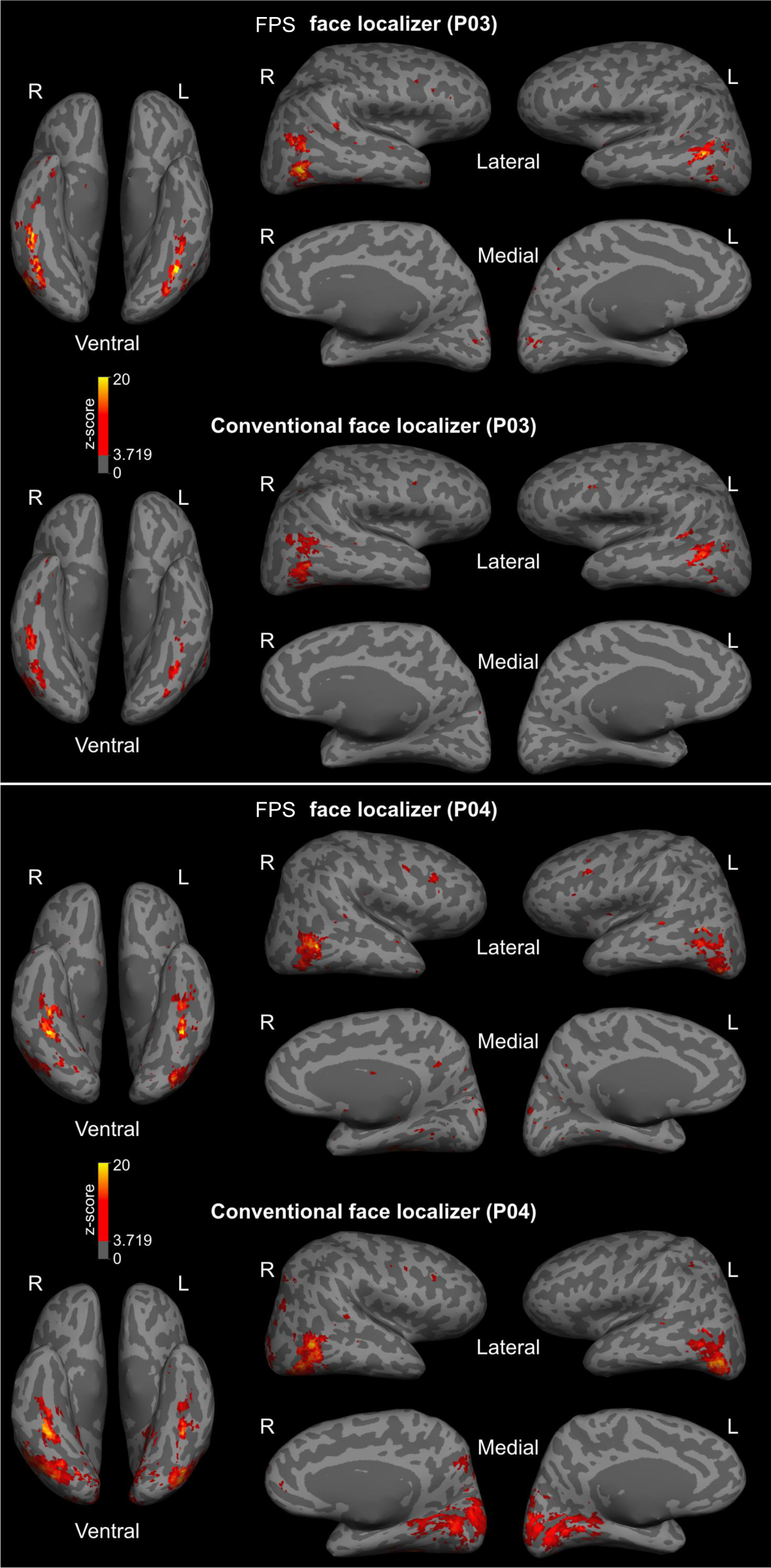
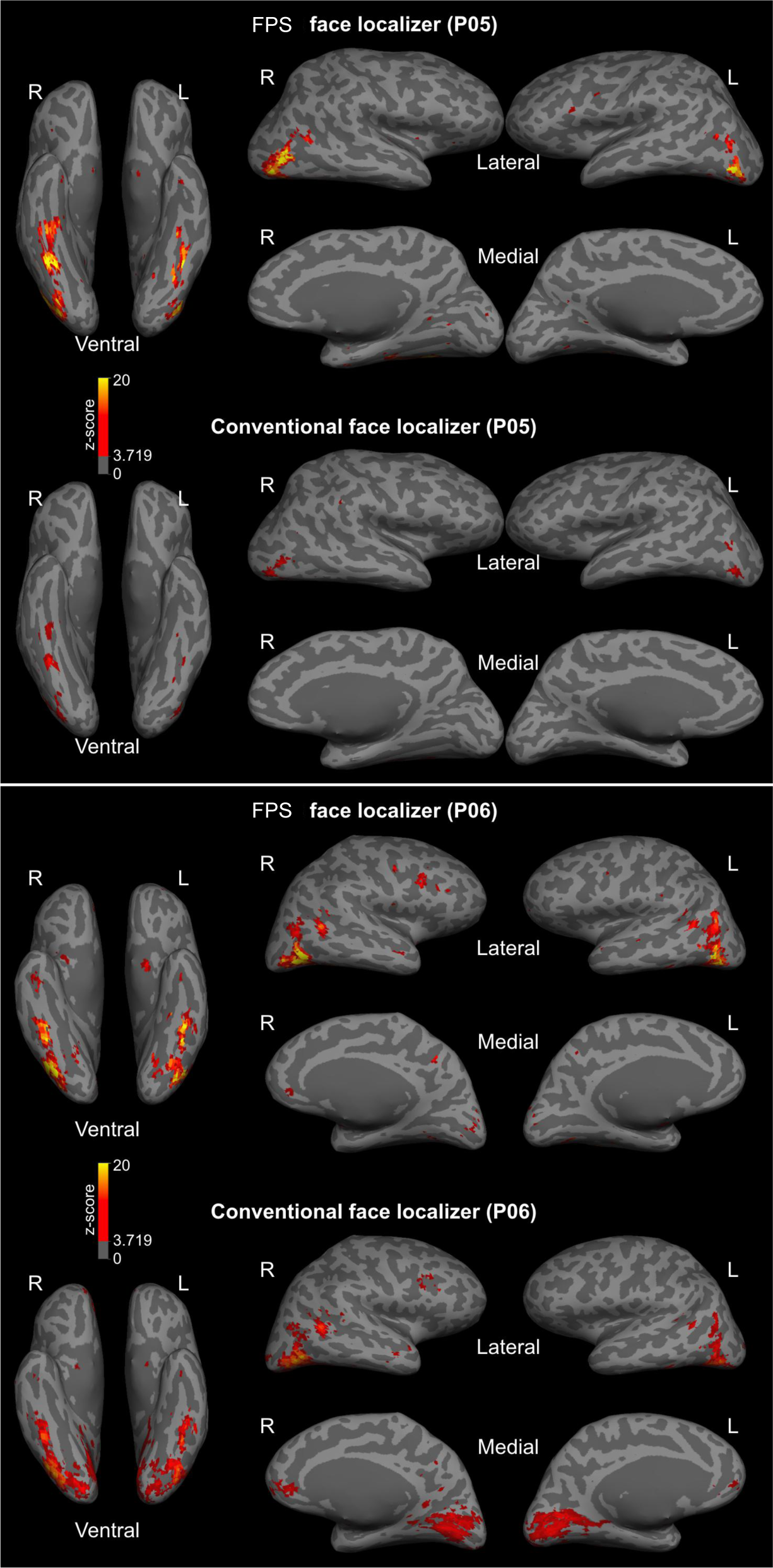
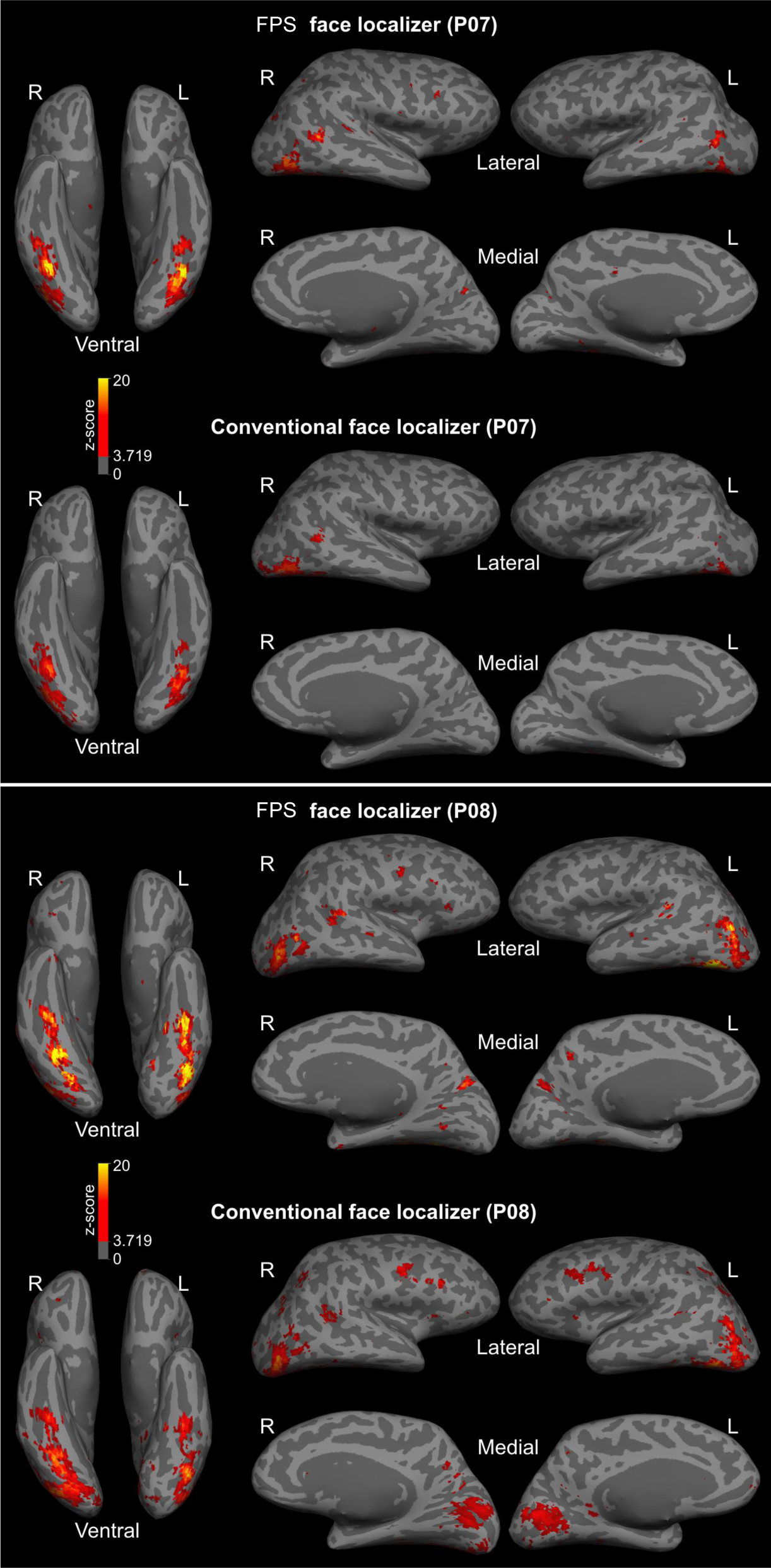
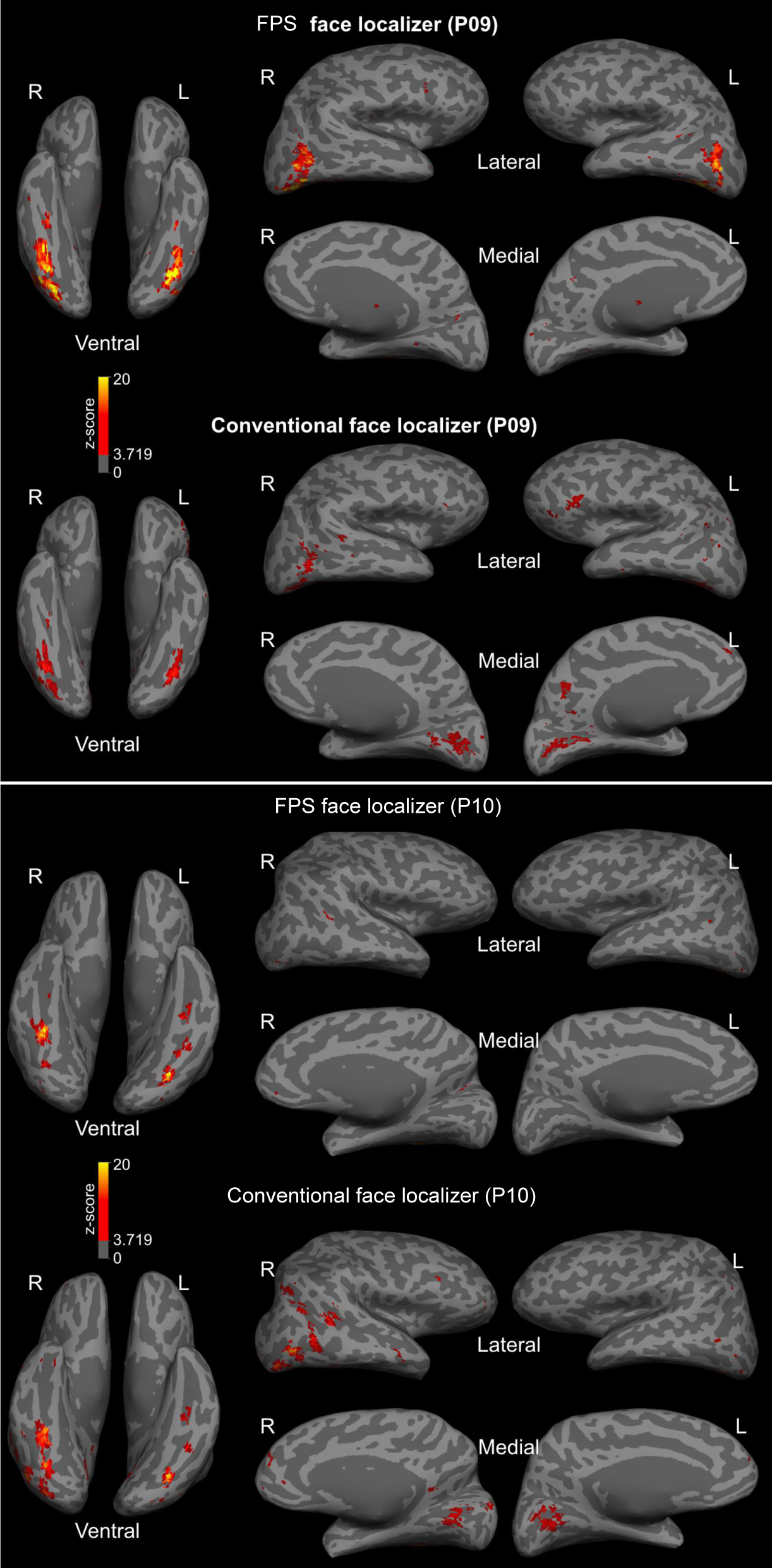
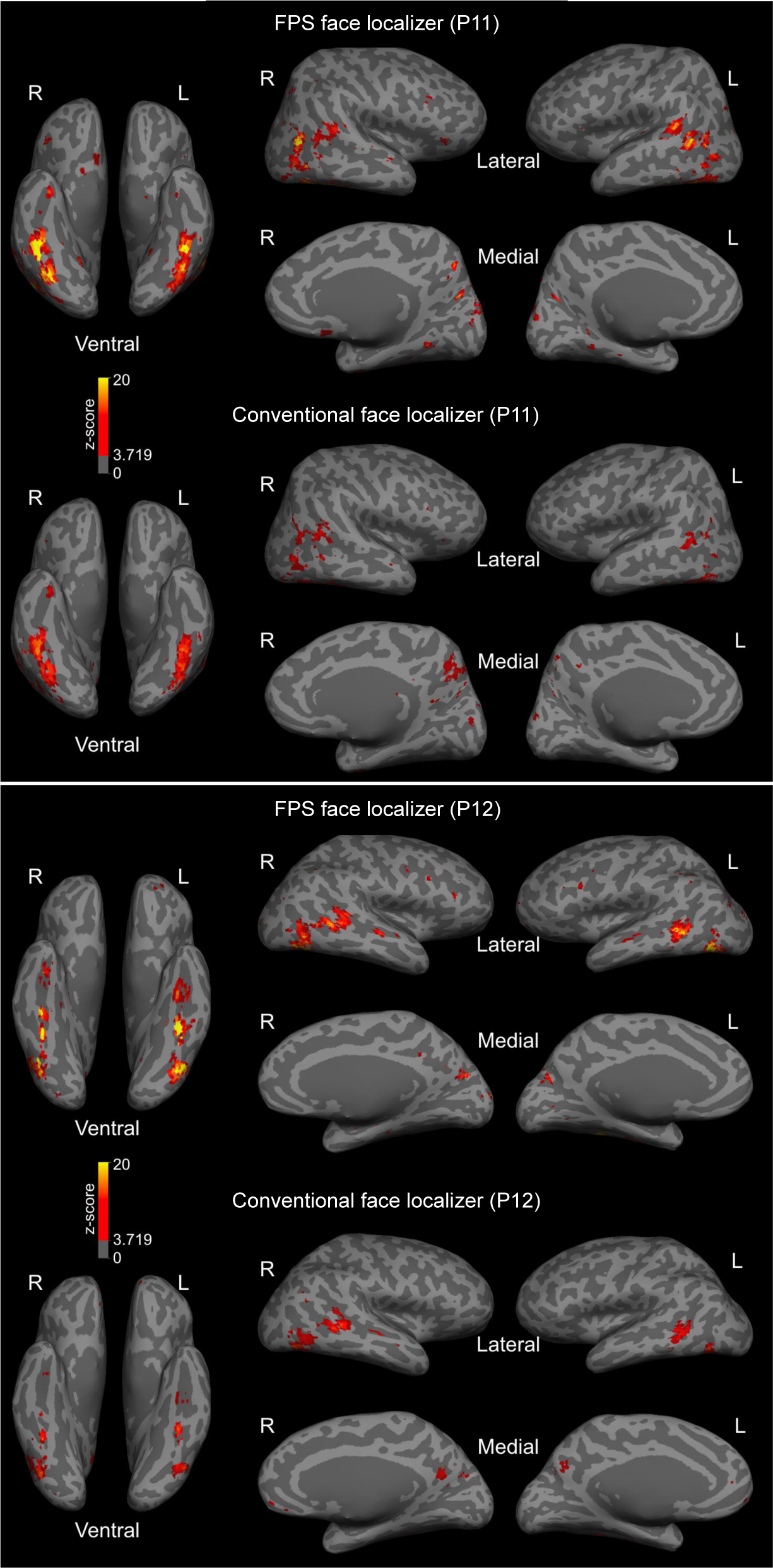
Face-selective areas in individual brains (P01 to P12) identified by the FPS-face localizer or by the Conventional face localizer (*p* < 0.0001, uncorrected). Related to Figure 4.

**Supplemental Fig. S6:**
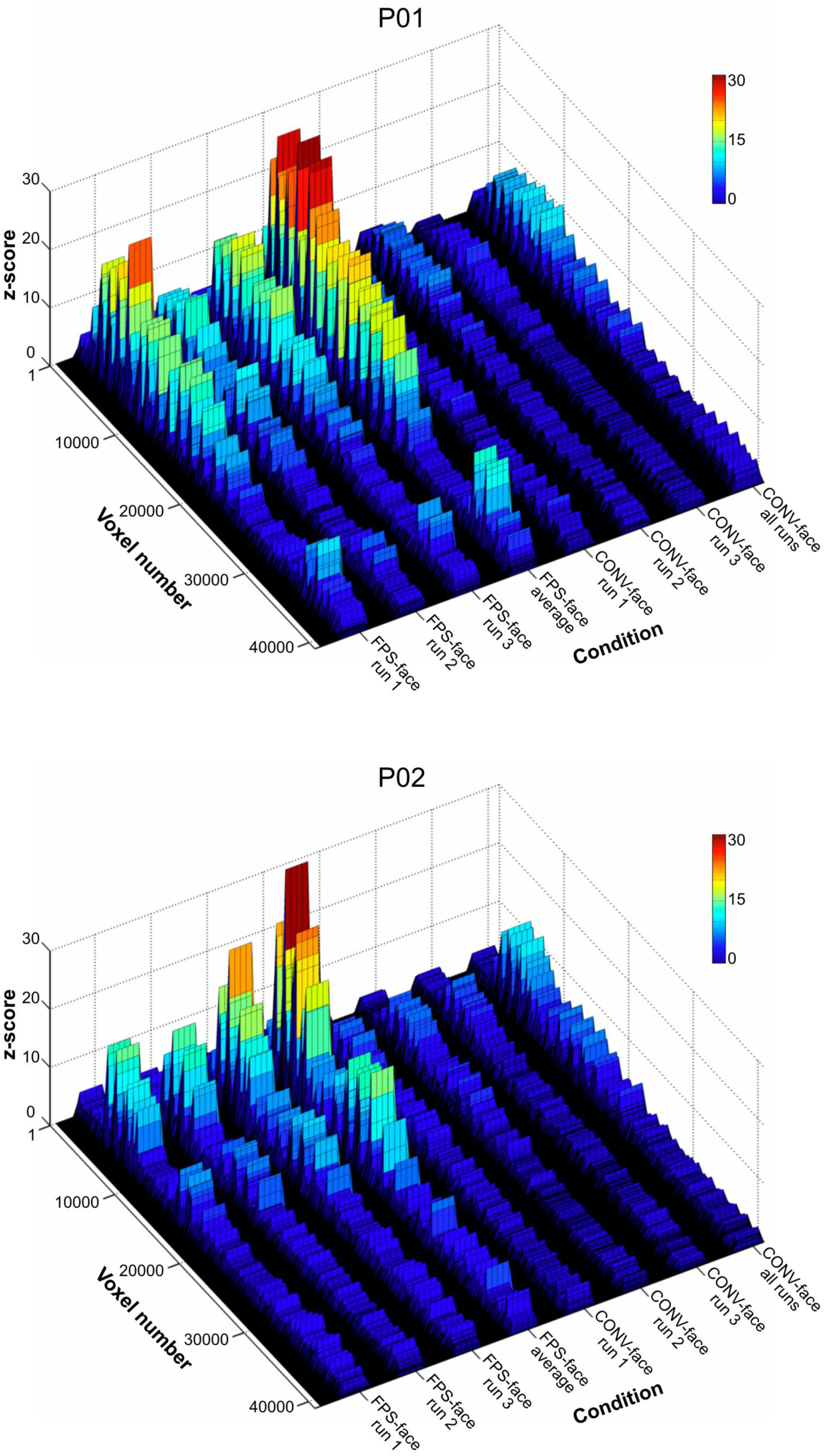
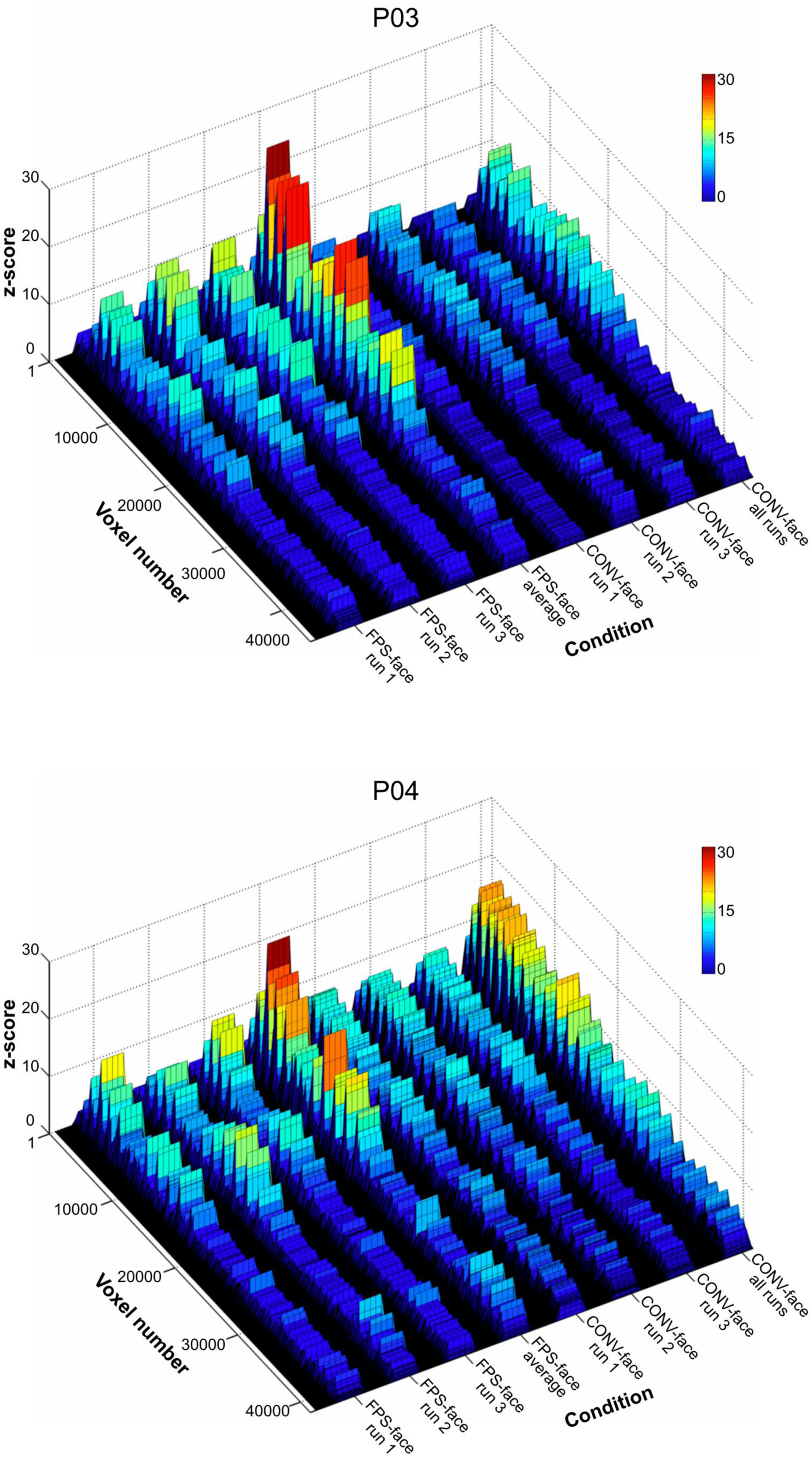
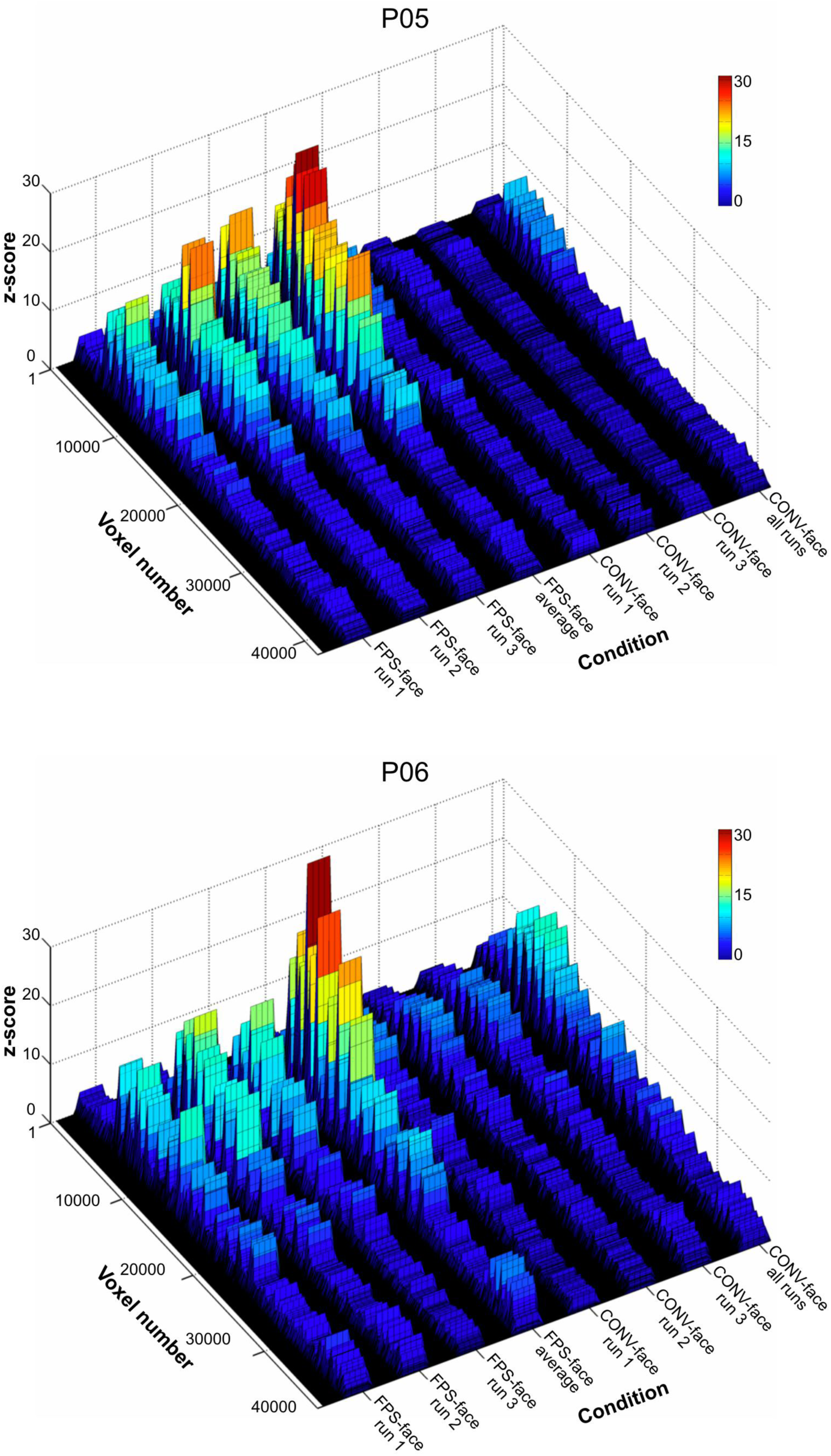
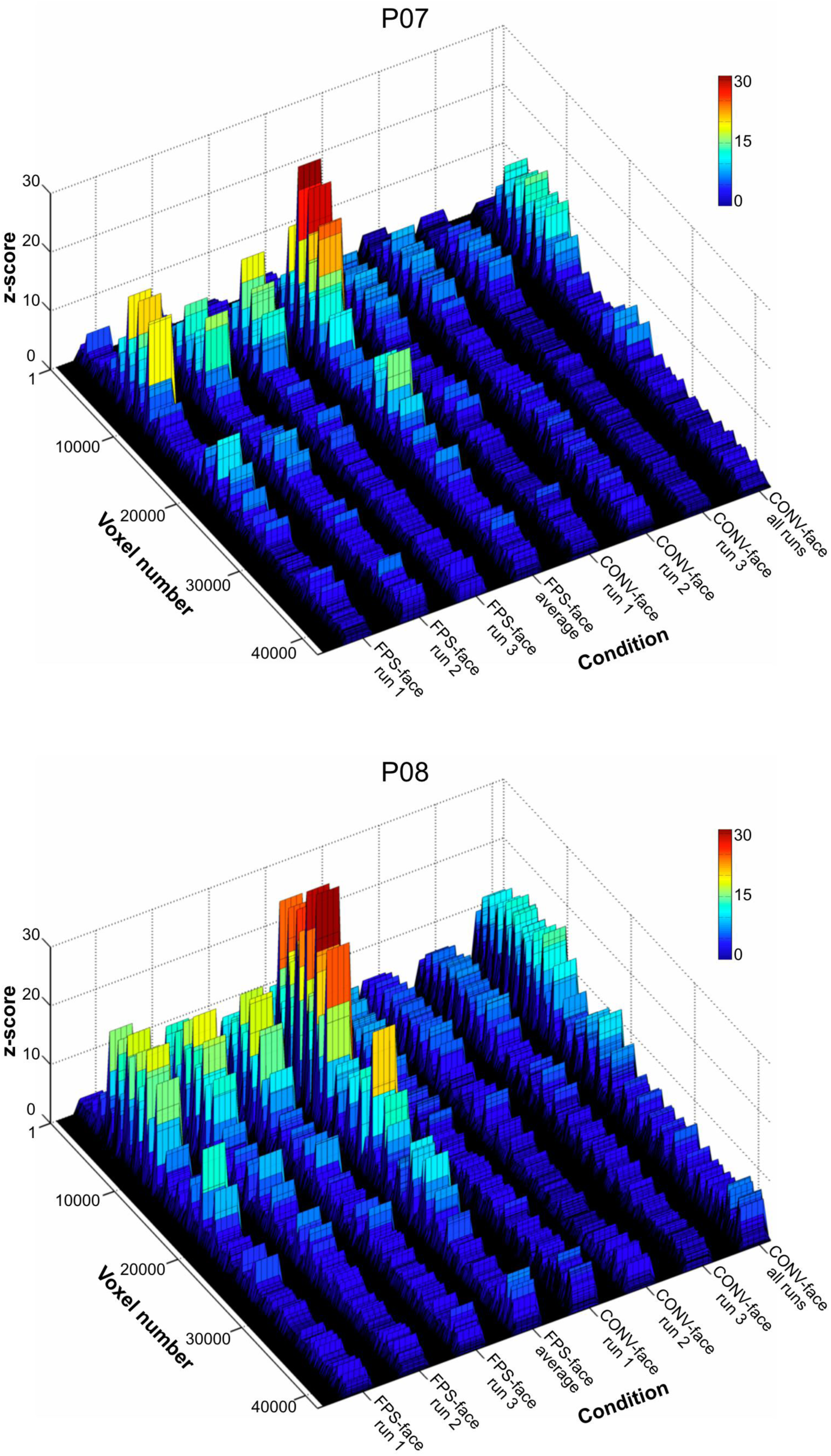
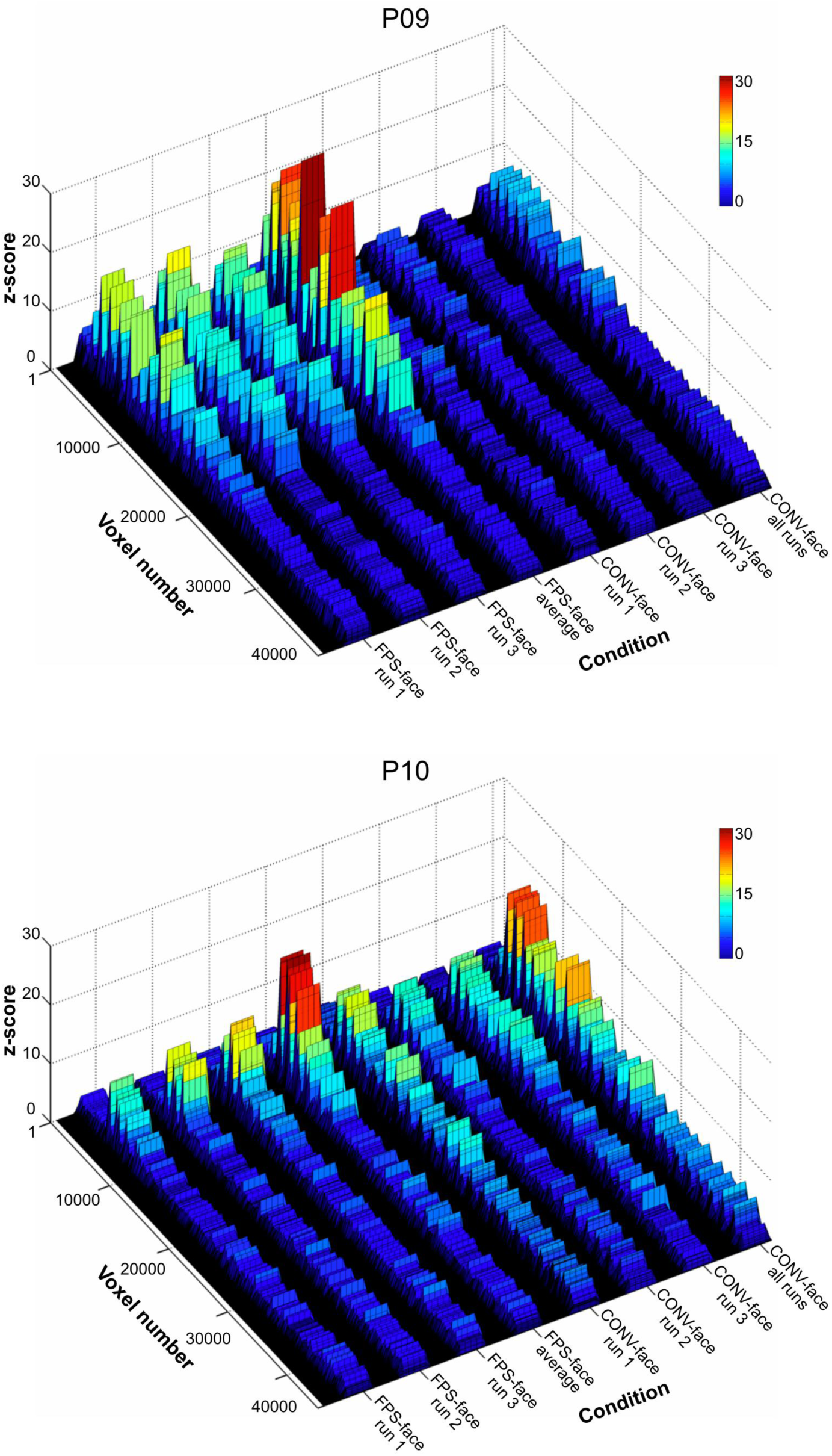
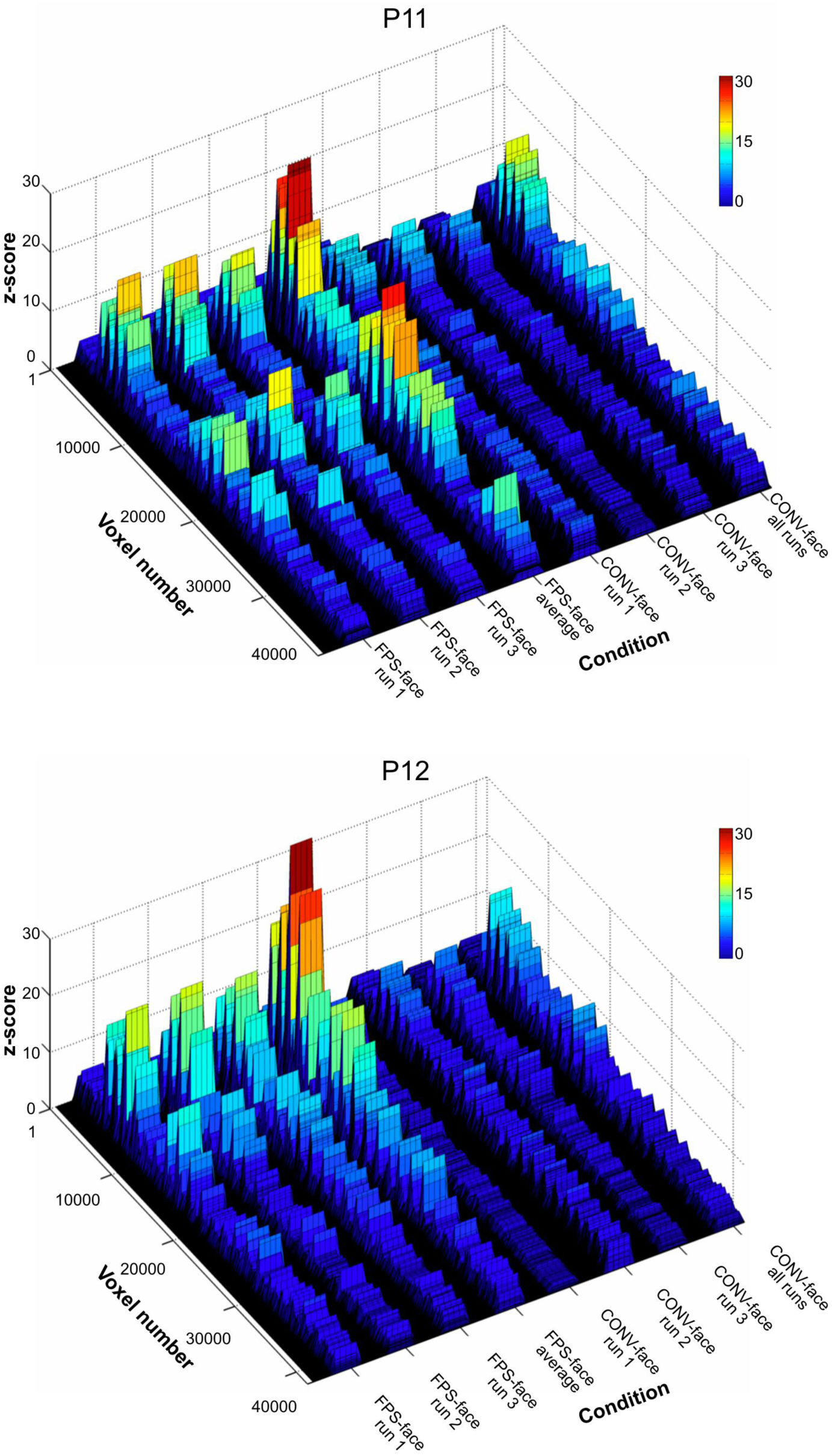
Face-selective neural response in individual runs and all runs combined in the FPS-face condition and CONV-face condition. Related to Figure 6.

**Supplemental Fig. S7:**
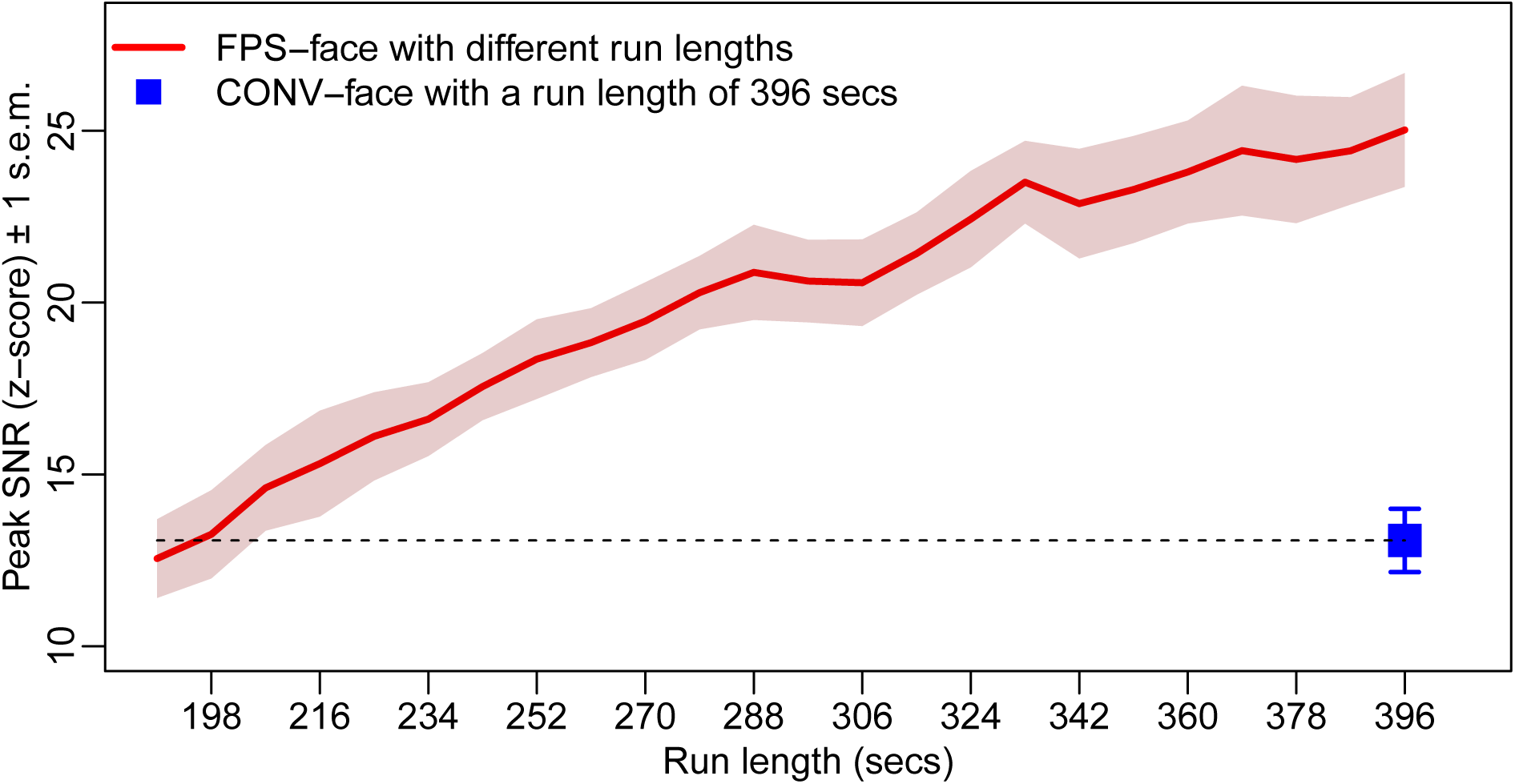
Peak SNR of the FPS-face condition with reduced scan duration. Related to discussion. Peak SNR of face-selective activity gradually decreases with reduced scanning duration. At a length 198 secs, which is half of the original scanning duration (396 secs), the peak SNR of the FPS-face condition has the same value as the peak SNR of the CONV-face condition with the original scanning duration (396 secs). Therefore, the FPS-face paradigm can effectively reduce the scanning time by a factor of two.

